# Self-replication of A*β*_42_ aggregates occurs on small and isolated fibril sites

**DOI:** 10.1101/2023.07.05.547777

**Authors:** Samo Curk, Johannes Krausser, Georg Meisl, Daan Frenkel, Sara Linse, Thomas C. T. Michaels, Tuomas P. J. Knowles, Anđela Šarić

## Abstract

Self-replication of amyloid fibrils via secondary nucleation is an intriguing physicochemical phenomenon in which existing fibrils catalyse the formation of their own copies. The molecular events behind this fibril surface-mediated process remain largely inaccessible to current structural and imaging techniques. Using statistical mechanics, computer modelling, and chemical kinetics, we show that the catalytic structure of the fibril surface can be inferred from the aggregation behaviour in the presence and absence of a fibril-binding inhibitor. We apply our approach to the case of Alzheimer’s A*β*_42_ amyloid fibrils formed in the presence of proSP-C Brichos inhibitors. We find that self-replication of A*β*_42_ fibrils occurs on small catalytic sites on the fibril surface, which are far apart from each other, and each of which can be covered by a single Brichos inhibitor.

---

The formation of amyloid fibrils is a ubiquitous form of protein self-assembly [1]. Despite their apparent simplicity – such fibrils are typically homomolecular and possess a quasi one-dimensional geometry – a number of distinct processes simultaneously participate in their formation. Fibril formation necessarily includes a nucleation step in which the initial fibrils form, followed by fibril elongation [2]. In addition, branching, lateral association, and self-replication processes such as fragmentation and secondary nucleation can be involved [1, 3–5]. Fibril self-replication by secondary nucleation in particular has emerged as a general feature of pathological protein self-assembly, observed in the context of many disorders [6]: haemoglobin fibrils involved in sickle cell anemia [7, 8], and amyloid fibrils associated with Alzheimer’s disease (Amyloid-*β* peptide, A*β*) [9, 10], type II diabetes (islet amyloid peptide, IAPP) [11–13], and Parkinson’s disease (*α*-synuclein) [14–16], are all able to make copies of themselves without any external energy input or complex cellular machinery. During secondary nucleation the surface of existing parent fibrils is able to catalyse the formation of daughter fibril nuclei. Due to this autocatalytic replication, which has been reported to be many orders of magnitude faster than nucleation not involving existing fibrils, the formation of new amyloid fibrils and the associated cytotoxicity becomes challenging to control [17, 18].

To date a number of successful experimental strategies have been reported to impede the self-replication of amyloid fibrils *in-vitro* [19–26]. In addition to emerging as a potential therapeutic strategy, the introduction of inhibitors into a self-replicating fibril system offers a controlled way of perturbing the system and gaining mechanistic insights. In this paper we use inhibitors, alongside a combination of statistical mechanics, chemical kinetics, and computer modelling, to show that kinetic signatures of amyloid aggregation in the presence of inhibitors can provide significant information about the catalytic structure of the fibril surface. We apply our approach to the case of the amyloid *β* peptide (A*β*_42_) from Alzheimer’s disease in the presence of proSP-C Brichos, a known inhibitor of secondary nucleation [27, 28]. A clear picture emerges from our study: A*β*_42_ secondary nucleation takes place on small and isolated catalytic sites, each of which can be completely inhibited by single Brichos molecules. We anticipate that the approach presented here can be used to interrogate nucleation and inhibition mechanisms behind self-replication of other amyloidogenic proteins, as well as of heterogeneous supramolecular catalysis more generally.

## Method of scaling exponents

Self-replication of amyloid fibrils via secondary nucleation has been shown to be a multi-step mechanism driven by the adsorption of monomeric amyloidogenic proteins onto the surface of amyloid fibrils [29–34]. It has been shown that the secondary nucleation rate in the absence of inhibitors, *r*, depends on the coverage of the fibril surface by monomeric protein *θ*_0_ as:

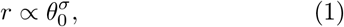

where *σ* is a scaling exponent [30, 33, 35]. This formula suggests that the nucleation process can be described as a rearrangement of *σ* surface-bound proteins into an oligomer of a critical size. This pre-equilibrium reaction step is then followed by a conformational conversion step from the surface-attached oligomer into a growth-competent *β*-sheet-rich fibrillar form (Fig. 1a, left). Using Langmuir adsorption model, the protein coverage can be expressed as:

**Figure 1.**
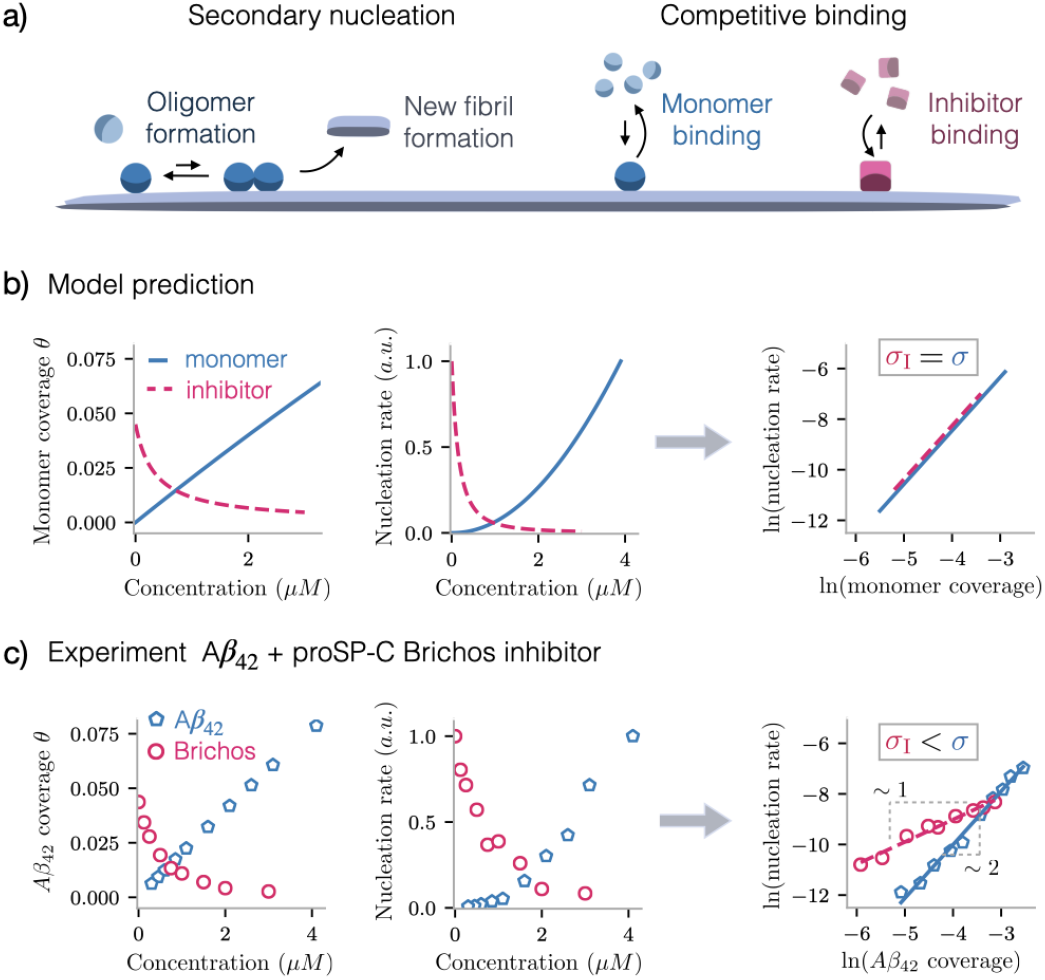
Fibril self-replication and its inhibition: a) Self-replication of amyloid fibrils by secondary nucleation involves binding of amyloidogenic protein monomers to the fibril surface, association of monomers into protein oligomers, and the catalysis of the oligomers into new daughter fibrils. The rate of this multi-step process is directly related to the coverage of the fibril surface by protein monomers. Nucleation of protein can be disrupted by inhibitors that decrease protein coverage by competing with monomers for the fibril surface. b) Langmuir model prediction for how protein coverage (left panel) and nucleation rate (middle panel) changes when modulating either monomer (blue solid line) or inhibitor (pink dashed line) concentration. Here, inhibitors only affect the protein coverage so the scaling exponents *σ, σ*_I_ that track the dependence of rate on coverage (defined in Eq. 4) should have the same value, regardless of whether protein coverage is modulated by changing protein or inhibitor concentration. c) In experiments, however, we observe starkly different behaviour; the values of scaling exponents differ significantly and the presence of inhibitors seems to increase the rate of nucleation at a given amount of protein coverage. Data for varying monomer is taken from [10] and data at varying inhibitor is same as in Fig. 4.

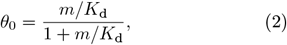

where *m* is the concentration of free monomers in solution, and *K*_d_ is the dissociation constant for the binding of monomers to the fibril surface. If we now introduce a fibril-binding inhibitor at concentration *I* into the system, the inhibitor will compete with monomers for space on the fibril surface and therefore decrease the protein surface coverage (Fig. 1a, right). This effect of a competitively binding inhibitor on the protein coverage can be expressed as:

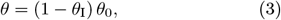

where *θ*_*I*_ is the inhibitor coverage of the fibril surface, and *θ* is the protein coverage that now depends on both protein and inhibitor concentration (see Supplementary Information for the derivation). Based on Eqs. 1-3, one would then expect that in the presence of inhibitors, the expression for the secondary nucleation rate (Eq. 1) would remain the same and only the protein coverage would be decreased.

To investigate how the presence of inhibitor influences the secondary rate, it is useful to define two scaling exponents that can be compared:

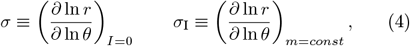

where *σ*, as already defined by Eq. 1, tracks the dependence of the secondary nucleation rate *r* on protein coverage *θ* in the absence of inhibitor. The second scaling exponent *σ*_I_ tracks the same dependence when inhibitor concentration is varied at constant monomer concentration. In a simple scenario where inhibitors displace monomers by competing with them anywhere on the surface (Figure 1a and Eqs. 1-3) one would expect the two scaling exponents to be equal *σ*_I_ = *σ* (Fig. 1b).

## A*β*_42_ aggregation in the presence of Brichos shows non-trivial inhibition behaviour

We now evaluate the scaling exponents *σ* and *σ*_*I*_ from *in-vitro* experimental data of A*β*_42_ aggregation kinetics in the presence of proSP-C Brichos, a molecular chaperone that binds to A*β* fibril surfaces and specifically inhibits secondary nucleation but does not bind A*β* monomers [27, 28, 36–38]. Per Eq. 4, we analyse two datasets. In the first dataset, the initial monomer concentration is kept constant and the inhibitor concentration is varied, allowing the evaluation of *σ*_I_ (Fig. S1). In the second dataset [33], the initial concentration of A*β*_42_ monomers is varied in the absence of any inhibitor, allowing the evaluation of *σ* (Fig. S2). The secondary nucleation rate (numerator in Eq. 4) can be directly extracted parameter-free from the time-dependence of aggregated fibril mass, as measured by ThT fluorescence (Eq. S.3). To evaluate the protein coverage (denominator in Eq. 4), we rely on the binding kinetics of A*β*_42_ monomer and proSP-C Brichos adsorption to A*β*_42_ fibril surface (Eqs. S.6-S.7), as measured by surface plasmon resonance [33, 37].

When changing the fibril coverage by varying the monomer concentration in the absence of inhibitor, we measure a constant scaling exponent *σ* = 2.3 ± 0.4. Surprisingly, when modulating the inhibitor concentration, the scaling exponent is much smaller, and equals unity (*σ*_I_ = 1.0 ± 0.1) (Fig. 1c). Interestingly, the measured disparity between scaling exponents (*σ*_I_ *< σ*) suggests that at a given fibril coverage by protein, the rate of secondary nucleation is higher in the presence of inhibitors than in their absence. We find these values of the scaling exponents to be well-bounded and robust (Fig. S3). The somewhat unintuitive result *σ* ≠ *σ*_*I*_ disagrees with the prediction from the ideal inhibitor binding model and points to a possibly non-trivial inhibition mechanism by which Brichos chaperones affect secondary nucleation.

In the following sections, we investigate the possible surface-based inhibition mechanisms of secondary nucleation and look for the origin of the scaling exponent behaviour *σ*_I_ = 1 *< σ*. Firstly, we use a particle-based computer model of secondary nucleation to investigate the effect of interprotein interactions on the scaling exponents (Fig. 2). Then, we develop a more general statistical-mechanics model of secondary nucleation and its inhibition that also takes into account the structure of the fibril surface (Fig. 3). Finally, based on the statistical mechanics model that best explains the experimental data, we build a kinetic model of amyloid aggregation in the presence of inhibitors and test it against time-dependent experimental data (Fig. 4). The summary of the results is given in Fig. 5.

**Figure 2.**
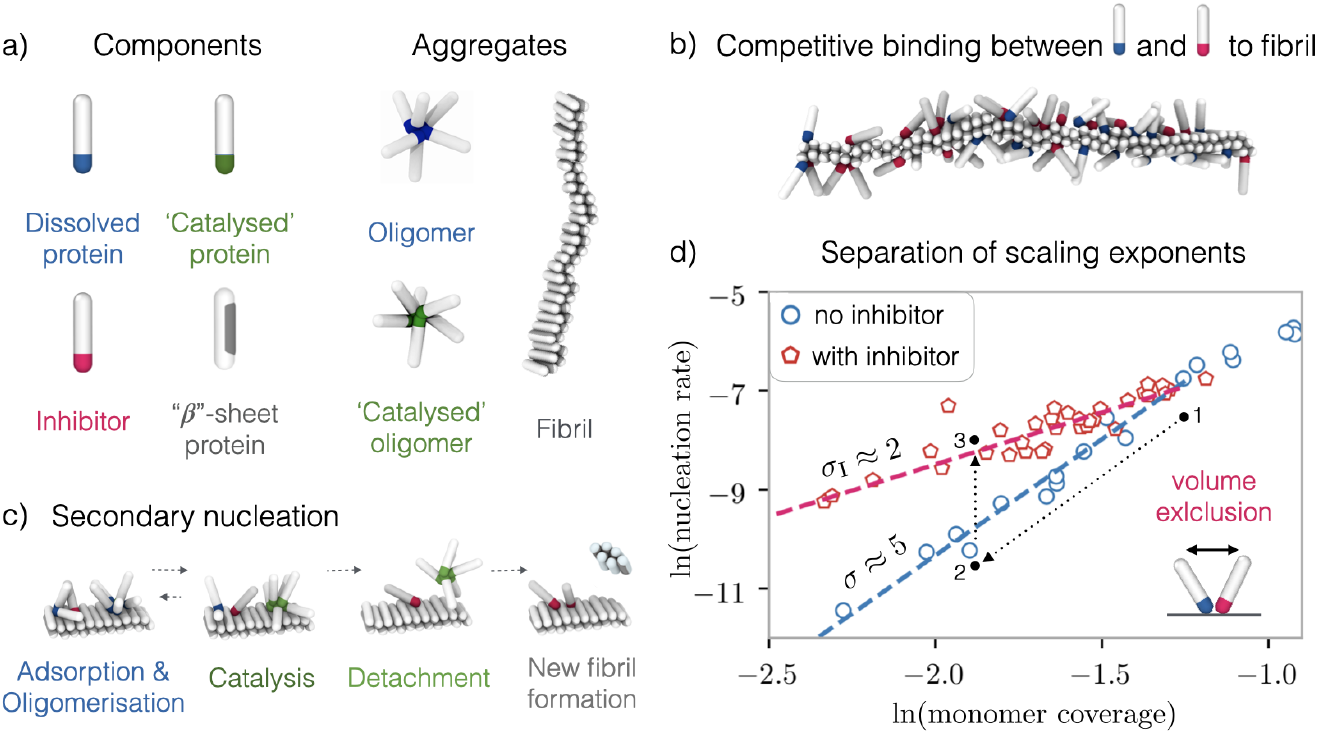
Particle-based model of secondary nucleation and its inhibition: a) Building blocks of the model are patchy spherocylinders with the capacity to self-assemble into oligomers and fibrilar structures. b) Inhibitors compete with protein monomers for common binding sites. c) The secondary nucleation pathway involves binding and oligomerisation of amyloid proteins on the fibril surface followed by an oligomer conversion to a catalysed, detachment-prone state, oligomer detachment, and finally a second conversion to a fibrillar form. d) Simulations reveal a stark mismatch of scaling exponents *σ* (slope of blue circles data) and *σ*_I_ (slope of red pentagons data) purely due to volume exclusion interaction between bound protein and inhibitor particles. Inhibitors act by lowering the protein coverage (path 1 *→* 2) through competitive binding but also promote micro-phase separation of proteins and inhibitors due to repulsive volume-exclusion interactions, leading to increased oligomerisation of protein and therefore enhancement of the nucleation rate (path 2 *→* 3).

**Figure 3.**
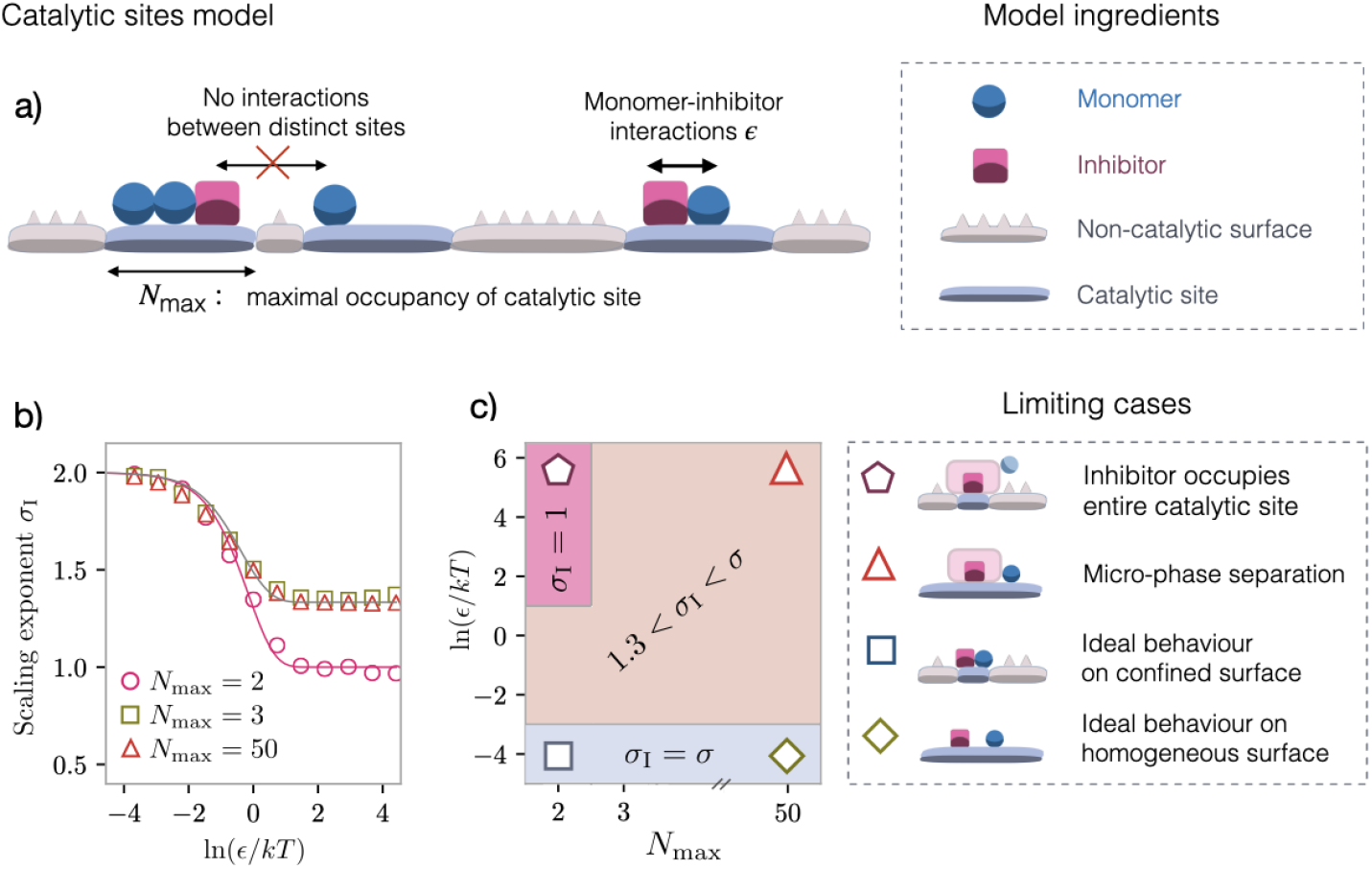
Independent catalytic sites model of the fibril surface: a) Model schematic: adsorption of protein monomers and inhibitors to the fibril surface occurs only on special catalytic sites that are separated by non-catalytic surface. Each catalytic site can hold up to *N*_max_ adsorbing particles. Neighbouring monomers and inhibitors bound to the same catalytic site interact with energy *ϵ* but there is no interaction of adsorbents between distinct catalytic sites. b, c) Value of the inhibition scaling exponent at varying strength of monomer-inhibitor repulsion energy and for different sizes of the catalytic surface, respectively. We recover the experimental value *σ*_I_ = 1 only in the limit where monomers and inhibitors are prevented from co-occupying the same catalytic site (*N*_max_ = 2, *ϵ → ∞*).

**Figure 4.**
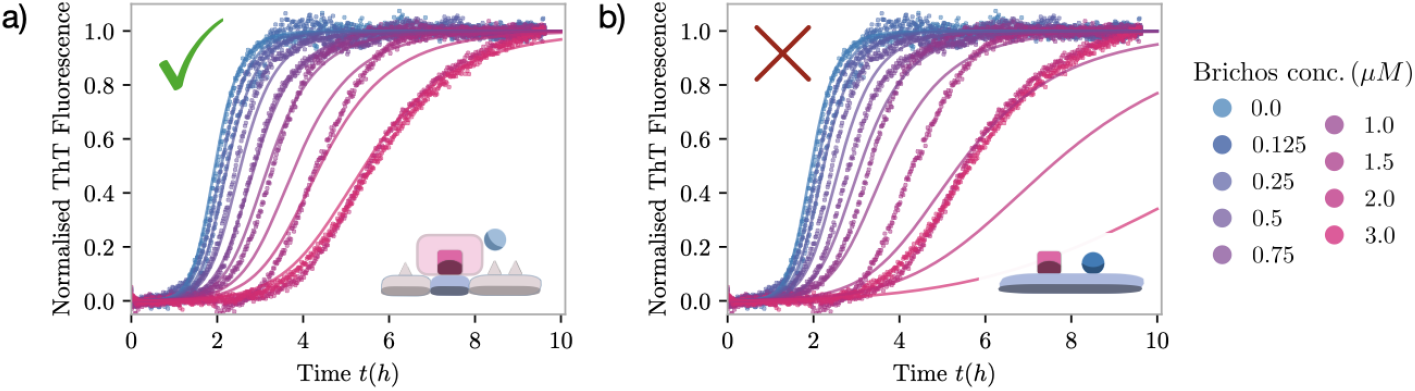
Kinetics of A*β*_42_ aggregation in the presence of surface-binding inhibitor: Time-dependence of fibril aggregation measured at 21°C in 20 mM sodium phosphate, 0.2 mM EDTA, pH 8.0, initial monomer concentration of 3 *µ*M, a low fibril seed concentration, and at varying concentration of proSP-C Brichos inhibitor. Datapoints represent normalised ThT fluorescence and the solid lines a global fit to the kinetic reaction schemes based on two limiting models of secondary nucleation and its inhibition. For model fit a) secondary nucleation takes place on small and distant catalytic sites, that can each be covered by a single inhibitor molecule. And for model fit b) inhibitors compete for the fibril surface according to Langmuir competitive binding kinetics.

**Figure 5.**
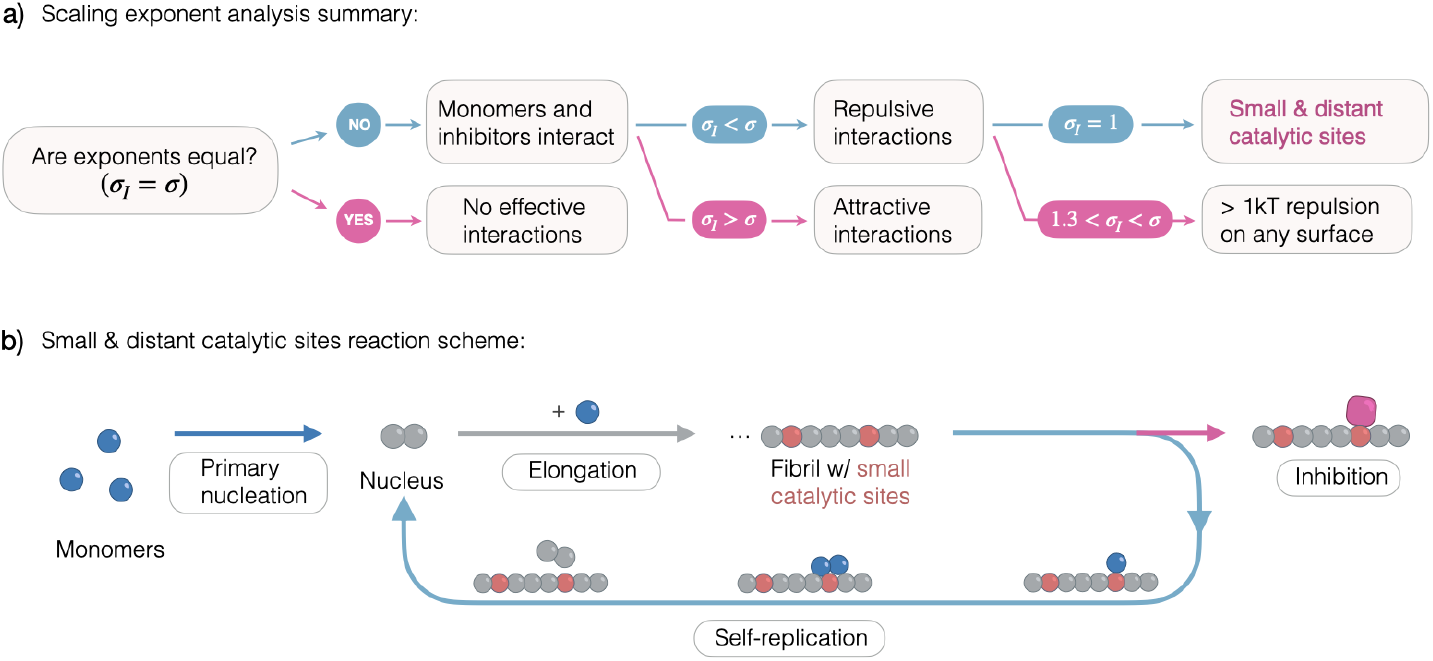
Summary of our analysis and the proposed model of A*β*_42_ secondary nucleation at small and isolated fibril sites. a) The method of scaling exponents (*σ, σ*_I_) allows us to extract information on the sign of interactions between fibril-adsorbed monomers and inhibitors, and in the special case of *σ*_I_ = 1, we additionally acquire information about the catalytic structure of the fibril surface. b) Our proposed kinetic reaction scheme for A*β*_42_ fibril aggregation in the presence of proSP-C Brichos. Fibril formation includes a primary nucleation step in which the initial fibrils form, followed by elongation of the fibril by addition of monomers to fibril ends. The surface of fibrils is able to catalyse the formation of new fibrils on special catalytic sites, leading to a fibril self-replication mechanism. This process of self-replication can be inhibited by Brichos inhibitors binding to catalytic sites, covering them whole.

## Particle-based model: monomer-inhibitor interactions influence *σ*_I_

The simple theoretical scenario for secondary nucleation and its inhibition presented in Fig. 1a neglects possible interactions between adsorbed particles. Realistically however, the few nanometers wide fibrils offer only a limited surface for protein and inhibitor binding, and one can imagine that non-ideal properties of adsorbed macro-molecules can alter the expected kinetic behaviour. To investigate the potential role of the macromolecular properties of the adsorbed molecules on secondary nucleation, we utilise a coarse-grained computer model of fibrillisation that describes all molecules as physical particles [30, 35, 39–43].

The amyloidogenic protein monomers are described as spherocylindrical particles (Fig. 2a) that can interconvert between three distinct conformational states: (i) the dissolved state which can transiently aggregate via non-specific interactions into unstructured oligomers, (ii) the *β*-sheet-prone state that readily assembles into compact elongated fibrillar structures, and (iii) the ’catalysed’ state on the fibril surface that stabilises a surface oligomer and enables its detachment from the surface. Energetically, the fibril-forming *β*-sheet-prone state is separated from the dissolved state by a large free-energy penalty. This creates a barrier for fibrillisation that can be effectively surpassed through a secondary nucleation process that involves the formation of an oligomer on the fibril surface, its conversion to the ’catalysed’ conformational state, detachment, and a final conversion to a fibril nucleus (see Fig. 2c), as explored in previous computational and experimental work on fibril self-replication [30, 35, 43].

We extend this scheme by introducing inhibitory molecules that bind to the fibril surface via the same adsorption mechanism as monomers. This results in a competition for the fibril surface between nucleation-able monomers and inert inhibitors (Fig. 2b). We performed grand-canonical Monte-Carlo simulations at different combinations of protein and inhibitor concentrations. We recorded the rates of fibril self-replication and protein coverage, just as in experiments, and evaluated the scaling exponents *σ* and *σ*_I_.

In the simulations, we observe a similarly stark disparity between the two scaling exponents *σ, σ*_I_ as in the experiment (Fig. 2d). Conveniently, simulations enable us to investigate the physical mechanisms behind this behaviour. We find that the scenario *σ*_I_ *< σ* is a result of two competing effects of inhibitors on secondary nucleation. Foremost, by competing for the same fibril surface, inhibitors substantially decrease the protein coverage and therefore decrease the rate of nucleation (see path 1 → 2 in Fig. 2d). However, at the same time, the repulsive volume-exclusion interaction between the macromolecular proteins and inhibitors effectively promotes micro-phase separation between the two bound species. This means that at a constant protein coverage, adding inhibitors promotes oligomerisation of protein on the fibril surface and increases the rate of nucleation (path 2 → 3 in Fig. 2d) relative to what we would expect from ideal Langmuir competitive binding. Although the presence of inhibitors leads to an overall slowing down of the nucleation process (Fig. S4), their ability to displace proteins from the fibril surface is partially offset by inhibitors’ influence on oligomerisation. To further confirm this, we performed simulations with appreciable attractive tip-to-tip interactions between inhibitors and monomers – in addition to their hard-core repulsion – and indeed, we measured *σ*_I_ *> σ* (Fig. S4). However, in our simulations, under no conditions could we recover the values of scaling exponents that emerged in experiment: while a repulsive interaction between adsorbed inhibitors and protein is sufficient to give an appreciable separation of scaling exponents (*σ*_I_ *< σ*) it cannot quantitatively explain experimental scaling (*σ*_I_ = 1).

## Statistical-mechanics model: *σ*_I_ = 1 points to small and isolated secondary nucleation sites

Our particle-based computer model suggests that a comparison between the two scaling exponents *σ* and *σ*_*I*_ can serve as a readout of the interactions between fibril-bound molecules. To explore broader conditions under which we can recover the experimentally measured *σ*_I_ = 1 we develop a general statistical-mechanical representation of secondary nucleation and its inhibition.

To start, we define all fibrils as an ensemble of independent subsystems that are surrounded by a common protein-inhibitor solution. The surface of each independent fibril is able to accommodate up to *N*_max_ molecules such that any number of monomers (*N*_m_) and inhibitors (*N*_I_) can adsorb subject to a constraint *N*_m_ + *N*_I_ ≤ *N*_max_. The adsorption of both species and their clustering on a given fibril surface is governed by a grand-partition function *ξ*(*λ*_m_, *λ*_I_, *N*_max_, *T*), where *λ*_m_ and *λ*_I_ are the monomer and inhibitor thermodynamic activities, respectively, and *T* is the temperature. Generally, we have:

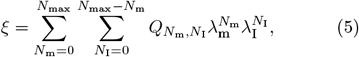

where 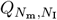 is the canonical partition function for the case with adsorbed *N*_m_ monomers and *N*_I_ inhibitors. The coverage of the fibril surface by the nucleating species *θ* and the rate of nucleation *r* can be extracted from the grand partition function as

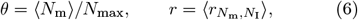

where ⟨*N*_m_⟩, and 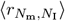 are the grand-canonical averages of the monomer occupancy, and per-occupancy rate of nucleation, respectively. The per-occupancy rate is in turn given by a canonical ensemble average over all different configurations *i* that *N*_m_ monomers and *N*_I_ inhibitors can arrange into:

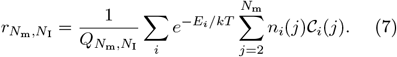

Here, *E*_*i*_ is the energy of a given configuration, *k* the Boltzmann constant, *n*_*i*_(*j*) the protein cluster distribution at configuration *i* (with *j* the size of the cluster), and 𝒞_*i*_(*j*) the rate of conformational conversion of a protein cluster of size *j* at the configuration *i*. This general form of the nucleation rate accounts for the possibility that protein oligomers of different sizes and shapes might get catalysed at different rates [44–47]. Combining the formulas for the average rate and coverage (Eq. 6), we can evaluate the scaling exponents *σ* and *σ*_I_ by enforcing the constraints *λ*_I_ = 0 or *λ*_m_ = *const*., respectively.

Remarkably, regardless of the potentially complicated details of the nucleating system and the influence of inhibitor, we can find a general condition under which *σ*_I_ = 1, which can be mathematically expressed as:

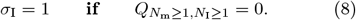

This vanishing partition function means that monomers and inhibitors cannot co-occupy the same space for *σ*_I_ = 1. Taking this consideration into account, the no co-occupancy 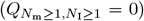 criterion can only be satisfied if a given fibril surface is small enough that a single volume-excluding inhibitor completely covers it and there is no space for a protein to bind at the same time. A more physical interpretation is that the fibril surface is divided into one or more catalytically active sites, where monomers and inhibitors can adsorb, and inactive surface that separates the active sites. Such catalytic sites have a small maximum occupancy *N*_max_, and the criterion of no monomer-inhibitor co-occupancy can therefore be easily satisfied. Hence our ensemble of independent subsystems refers to an ensemble of catalytic sites. To be considered independent, such sites need to be far apart from one another such that molecules adsorbed onto them cannot cross-interact.

Let us now consider a simplified case that emerged from the above general statistical-mechanics model, for which we can more easily evaluate the scaling exponents: a one-dimensional fibril covered with sparse catalytic sites. Monomers and inhibitors can bind the catalytic sites and interact with each other with an energy *ϵ* (Fig. 3a). Without loss of generality, we specify the size of the nucleation-prone oligomer to be 2, which sets the conformational conversion rate of protein clusters of size *j* to scale as 𝒞_*i*_(*j*) ∝ *j* − 1. Consequently, the scaling exponent in the absence of inhibitor and monomer-monomer interactions is consistent (*σ* = 2) for all *N*_max_. We then solve this simplified model for the scaling exponent *σ*_I_ at different values of monomer-inhibitor interaction energy *ϵ* and for different values of *N*_max_.

In the ideal Langmuir competitive binding regime (*ϵ* → 0), we find *σ*_I_ = *σ* = 2, as expected from Fig. 1, regardless of the maximal occupancy of the catalytic site (Fig. 3b, top-left). However, as soon as monomers and inhibitors interact with a non-vanishing repulsive interaction, *ϵ >* 0, *σ*_I_ acquires values appreciably below *σ* for *ϵ* ∼ *kT* and then reaches a constant value at higher repulsion energies (Fig. 3b, middle-right). The value to which *σ*_I_ saturates depends on the maximal occupancy of the catalytic site. For instance, for *N*_max_ = 50, which mimics a scenario where the entire fibril surface contributes to secondary nucleation akin to a homogeneous fibril, the inhibition scaling exponent saturates to about *σ*_I_ = 1.3 (red triangles in Fig. 3b). The value of *σ*_I_ remains approximately the same as long as *N*_max_ ≥ 3 (Fig. 3b). However, for *N*_max_ = 2 (pink circles in Fig. 3b), and only then, do we find *σ*_I_ = 1. In other words, the experimentally observed *σ*_*I*_ can be recovered only if the catalytic site is small (*N*_max_ = 2) and if the inhibitor and monomer interact strongly unfavourably with each other (*ϵ >>* 0). This strong unfavourable interaction in practice means that a monomer and inhibitor cannot occupy the same binding site, which can be satisfied if the monomer and inhibitor cannot overlap on a small binding site due to volume-exclusion. No other repulsive interaction is required.

This result confirms the no co-occupancy criterion of Eq. 8. It is important to notice that the uncertainty in the experimentally measured *σ*_I_ is much smaller than the difference between *σ*_I_(*N*_max_ = 2, *ϵ* → ∞), where co-occupancy is prohibited, and *σ*_I_(*N*_max_ = 3, *ϵ* → ∞), where co-occupancy is possible. For completeness, we also considered alternative scenarios for secondary nucleation to the one outlined in Eq. 7. In all the cases however we found that *σ*_I_ = 1 requires small catalytic sites and no co-occupancy between monomer and inhibitor (see Supplementary Information).

## Small fibril sites model explains A*β*_42_ aggregation kinetics in the presence of proSP-C Brichos

Let us now test the mechanistic picture of the fibril catalytic surface against time-dependent experimental data. To do so, we built a kinetic model of amyloid aggregation that incorporates small fibril sites and test its predictions against the time-dependent A*β*_42_ aggregation data.

Following previous work, we describe the aggregation kinetics of *Aβ*_42_ by combining primary nucleation, self-replication by secondary nucleation and fibril elongation into a single kinetic model that describes the number (*P*) and mass (*M*) concentrations of fibril aggregates as a function of time (*t*) [10, 29, 33, 48]:

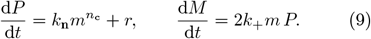

Here, *n*_c_ is the reaction order for primary nucleation, *k*_n_ and *k*_+_ are the rate constants for primary nucleation and elongation, respectively, and *r* is the rate of secondary nucleation. We model secondary nucleation and its inhibition by Brichos as:

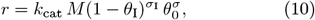

where *k*_cat_ is a catalysis rate constant for converting a surface oligomer into a new fibril nucleus, *θ*_*I*_ is the coverage of catalytic surface by Brichos, and *θ*_0_ the coverage by protein in absence of Brichos, given by the Langmuir isotherm of Eq. 2. Importantly, Brichos binding to the fibril occurs on a slow timescale, comparable to that of fibril aggregation [27, 37, 46]. Hence, unlike *θ*_0_, Brichos coverage cannot be captured by a pre-equilibrium expression. Rather, it evolves in time according to:

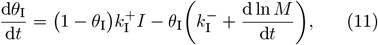

where 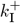 and 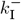 are the Brichos association and dissociation rate constant, and the term (d ln *M/*d*t*) signifies that as the amount of fibril surface increases through elongation, the proportion of the surface covered by inhibitor decreases. See Supplementary Information for a full derivation of the rate laws.

Depending on the chosen value of the inhibitor scaling exponent *σ*_I_, the same rate formula of Eq. 10 describes both the nucleation-inhibition scenario of Eqs. 1–3 (*σ*_I_ = *σ*), and the small fibril sites scenario that we derived from the general statistical mechanics model (*σ*_I_ = 1 *< σ*). To compare which model of the fibril surface and inhibition matches better with experiment, we numerically solve Eqs. 9–11 and fit the evolution of fibril mass *M* (*t*) against experimental data of A*β*_42_ aggregation at different concentrations (*I*) of proSP-C Brichos. We perform the fits in a global manner, where the same set of rate laws and a common set of kinetic parameters attempts to describe data at all inhibitor concentrations. The global parameters we fit are *k*_cat_ and the initial amount of fibril seed (*M*(*t* = 0)). All other parameters (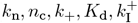, and 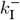) are taken from the literature [33, 37].

As shown in Fig. 4, both models fit the experimental aggregation data equally well in the absence of inhibitor where the value of *σ*_I_ does not play a role (left-most curve *I* = 0 in Fig. 4). However, at increasing concentrations of Brichos, it becomes clear that the model with small and independent secondary nucleation sites that can be covered by single inhibitors (Fig. 4a) outperforms the model where ideal inhibitors bind according to the Langmuir adsorption model (Fig. 4b).

Crucially, the comparison between the two models showcases that the inhibition mechanism and the associated value of *σ*_I_ greatly influence the aggregation kinetics. Moreover, it demonstrates that our analysis workflow - the extraction of experimental scaling exponents in the presence and absence of inhibitors - provides a method for finding molecular mechanisms and the corresponding rate laws.

## Discussion and Conclusions

In this paper, we have demonstrated how inhibitory molecules, beyond their therapeutic promise, can serve as a powerful tool for identifying mechanistic details underlying protein aggregation phenomena. In particular, the use of inhibitory molecules within statistical-mechanics and chemical kinetics frameworks revealed structural heterogeneity of *Aβ*_42_ fibril surface, where only specific and small sites on the fibril surface are able to catalyse the formation of new fibrils. Each such catalytic site is likely completely inhibited by a single proSP-C Brichos complex. Our model also suggests that these catalytic sites need to be substantially distant from one another such that the contact between bound inhibitors and proteins is prevented. Figure 5 presents the summary of our results.

The above requirements suggest a lower bound on the spacing between catalytic sites that we estimate to be at least the size of Brichos (∼ 15 nm [49, 50]). The distance between secondary nucleation sites is thus much larger than the repeat distance of A*β* monomers along the fibril axis, which is only a few Å [51]. While we should expect non-homogeneity of the fibril structure on the length scale of monomer spacing, we find the surface heterogeneity on a length scale that is larger than *>*∼ 10 nm surprising, especially given the universally homomolecular makeup of the fibrils [52].

There is, however, a body of evidence supporting the scenario of surface heterogeneity. Surface plasmon resonance measurements of Brichos binding to the fibril surface suggest two binding modes [37]. While this behaviour could arrise due to co-existence of strong specific and much weaker non-specific interactions with the homogeneous fibril surface, it could also indicate two types of binding sites. Additionally, electron-microscopy images of immunogold-labelled Brichos bound to the fibril surface suggest sparse binding sites [27, 36]. Furthermore, a recent co-aggregation study of A*β*_42_ and S100A9 proteins shows S100A9 fibrils binding to the surface of A*β* fibrils at a sub-stoichiometric ratio of one S100A9 fibril per about 300 fibrillar A*β* monomers while completely abolishing secondary nucleation [53]. The same study reports a smallest distance of ∼ 40 nm between surface-bound S100A9 fibrils, corroborating our model.

Our analysis cannot reveal the origin of secondary nucleation sites, but it can be hypothesised that such sites are not periodic, as otherwise they would be visible from structural data. They also do not seem to be correlated with morphological features of the fibril, such as the fibril local curvature or the relative intensity of ThT fluorescence, as recently reported [54]. Instead, it is tempting to speculate that catalytic sites might originate on the level of higher-order fibrillar assembly, such as bundling, braiding or coiling of filaments into mature fibril structures. This would however imply that secondary nucleation sites form on a timescale related to inter-filament kinetics, which is expected to be much slower than the timescale observed for secondary nucleation. Alternatively, random point defects arising from missing or misaligned fibrillar monomers could occur anywhere on the fibril surface due to thermal fluctuations. These defects would scale in number with the amount of aggregated fibril mass, consistent with Eq. 10, and their presence could expose the fibril core for motifs not present on the fibril surface [55, 56].

Importantly, the realisation that the fibril surface might be structurally heterogeneous has implications for structure-based rational drug design, as most discovery methods focus on structural information averaged along the fibril axis. Similarly, antibodies designed to target epitopes exposed on the fibril surface usually bind the fibril with very high affinity but do not seem to act as particularly good inhibitors of secondary nucleation [25, 26]. This indicates that such antibodies, including aducanumab that has recently been put in the spotlight in the context of Alzheimers’s disease treatment [57, 58], likely bind to non-catalytic parts of the fibril surface, hence interfering with secondary nucleation only indirectly. We expect that future work will provide more details into the nature of catalytic sites both via structural methods, as well as by measuring their persistence over multiple secondary nucleation events or their abundance at varying environmental conditions.

## Acknowledgments

We acknowledge support from the ERASMUS programme and the UCL Institute for the Physics of living Systems (S.C., T.C.T.M.), the Biotechnology and Biological Sciences Research Council (T.P.J.K.), the Engineering and Physical Sciences Research Council (D.F.), the European Research Council (T.P.J.K., S.L., D.F, and A.Š.), the Frances and Augustus Newman Foundation (T.P.J.K.), the Academy of Medical Sciences and Wellcome Trust (A.Š.), and the Royal Society (S.C. and A.Š.).

