## Supplementary Information for "Self-replication of A*β*_42_ aggregates occurs on small and isolated fibril sites"

**This PDF file includes:**

Figs. S1 to S5

### 1. Extraction of scaling exponents

The scaling exponents $\sigma$ and $\sigma_\mathrm{I}$ track the dependence of the rate of secondary nucleation ($r$) on the protein coverage of the fibril surface ($\theta$) when modulating protein concentration ($m$) or inhibitor concentration ($I$), respectively. In kinetic aggregation experiments, both $\theta$ and $r$ are not directly measurable so we need to evaluate them from other known variables. We use the following assumptions about aggregation and binding kinetics when performing our analysis:

1. Protein mass is mostly distributed between dissolved monomeric form, and the insoluble aggregated fibril form yielding the conservation law $m(t) + M(t) = m_\mathrm{tot}$, where $M$ is the monomer equivalent concentration of aggregated protein, and $m_{tot}$ is the total protein concentration. This has been reported to be correct to within 1% accuracy for A$\beta_{42}$ meaning that the contributions to protein mass from fibril-bound monomers and oligomers in solution is minimal [1].
2. The elongation of fibrils occurs by attachment of free monomers to fibril ends such that $\mathrm{d}M/\mathrm{d}t \propto m$ [2].
3. The measured intensity of the Thioflavin T (ThT) amyloid dye signal is directly proportional to $M$ [3].
4. The protein and inhibitor binding kinetics can be described by a competitive binding model where protein monomers and inhibitors compete for common fibrillar binding sites. See section 2.4 for a derivation of this.

**1.1. Evaluating the nucleation rate.** The overall rate of nucleation $r_\mathrm{total}$ is defined as:

$$r_\mathrm{total} \equiv \frac{\mathrm{d}P(t)}{\mathrm{d}t}, \tag{S.1}$$

where $P$ is the concentration of fibrils. As per assumption 2 we model fibril elongation as:

$$\frac{\mathrm{d}M(t)}{\mathrm{d}t} = 2k_+ P(t)\, m(t), \tag{S.2}$$

where $k_+$ is the elongation rate constant, and the factor 2 comes from free monomers being able to bind to both ends of the fibril. Now combining the assumptions 1 and 3 such that $M/m_{tot} \equiv \tilde{\mathcal{M}}$ is proportional to the normalised ThT signal with equations S.1 and S.2, we can express the overall rate of nucleation as:

$$r_\mathrm{total} = \frac{-1}{2k_+}\frac{\mathrm{d}^2}{\mathrm{d}t^2}\ln\left[1 - \tilde{\mathcal{M}}(t)\right]. \tag{S.3}$$

We see that the rate of nucleation can be therefore evaluated directly from the time-dependence of the normalised ThT signal. The secondary nucleation rate is then given by Eq. S.3 in the regime of non-negligible fibril mass where secondary nucleation rate is expected to dominate over the rate of primary nucleation ($r \approx r_\mathrm{total}$).

**1.2. Evaluation of coverages.** To evaluate the monomer ($\theta$) and inhibitor ($\theta_\mathrm{I}$) coverages at every time-point we treat the binding kinetics under the assumption 4. The competitive binding to the fibril surface can be expressed as:

$$\frac{\mathrm{d}M_\mathrm{m}(t)}{\mathrm{d}t} = k_\mathrm{m}^+ M_\mathrm{f}(t)\, m(t) - k_\mathrm{m}^- M_\mathrm{m}(t) \tag{S.4}$$

$$\frac{\mathrm{d}M_\mathrm{I}(t)}{\mathrm{d}t} = k_\mathrm{I}^+ M_\mathrm{f}(t)\, I(t) - k_\mathrm{I}^- M_\mathrm{I}(t), \tag{S.5}$$

where $M_\mathrm{m}$ is the concentration of aggregated (i.e. fibrillar) protein mass that is covered by monomers, $M_\mathrm{I}$ concentration of fibrillar protein covered by inhibitors, and $M_\mathrm{f}$ concentration of aggregated protein that is free from any adsorbate, with $k_\mathrm{m}^+$ ($k_\mathrm{I}^+$), and $k_\mathrm{m}^-$ ($k_\mathrm{I}^-$) the protein (inhibitor) adsorption and desorption rate constants, respectively. At any time, the fibril mass is conserved as: $M(t) = M_\mathrm{f}(t) + M_\mathrm{m}(t) + M_\mathrm{I}(t)$.
The binding kinetics of protein adsorption operate on a much faster timescale than fibril elongation under quiescent conditions [4], so we can set Eq. S.4 to zero and rewrite it as:

$$\theta(t) = (1 - \theta_\mathrm{I}(t))\, \theta_0(t), \tag{S.6}$$

where $\theta \equiv M_\mathrm{m}/M$, $\theta_\mathrm{I} \equiv M_\mathrm{I}/M$ are the protein and inhibitor coverages of the fibrils surface, respectively, and $\theta_0 = \frac{m(t)/K_\mathrm{d}}{1+m(t)/K_\mathrm{d}}$ is the protein coverage in the absence of inhibitor, with $K_\mathrm{d} = k_\mathrm{m}^-/k_\mathrm{m}^+$ the dissociation constant for monomer binding.
Contrary to A$\beta_{42}$ binding kinetics, Brichos binding operates on a much slower timescale that is comparable to A$\beta$ fibril elongation. By dividing both sides of Eq. S.5 with $M(t)$, taking into account Eq. S.6, and using the product rule we can write the differential equation for inhibitor coverage as:

$$\frac{\mathrm{d}\theta_\mathrm{I}(t)}{\mathrm{d}t} = \left[(1 - \theta_\mathrm{I}(t))\,(1 - \theta_0(t))\right] k_\mathrm{I}^+ I(t) - \theta_\mathrm{I}(t)\left[k_\mathrm{I}^- + \frac{\mathrm{d}}{\mathrm{d}t}\ln\tilde{\mathcal{M}}(t)\right], \tag{S.7}$$

where the last term on the right-hand side of Eq. S.7 signifies that as the amount of fibril surface increases through elongation, the proportion of the surface covered by inhibitor drops. To evaluate the protein coverage, we therefore need to numerically solve the set of Eqs. S.6 and S.7 with $\tilde{\mathcal{M}}(t)$ taken directly from experimental measurements of aggregation kinetics, and $k_\mathrm{I}^+$, $k_\mathrm{I}^-$, and $K_\mathrm{d}$ taken from the literature [4, 5]. Note finally that under dilute monomer conditions $m \ll K_\mathrm{d}$, the term $\theta_0(t)$ vanishes the differential equation for the inhibitor coverage (Eq. S.7). In this case inhibitor binding kinetics can be considered independent of monomer binding and we get the expression Eq. (11) in the main text. We discuss this operation further in subsection 2.4

**1.3. Evaluating $\sigma_{\mathrm{I}}$ and $\sigma$.** For slow inhibitor binding kinetics the coverage of the fibril surface by inhibitor and protein adsorbates follows with considerable delay the conditions imposed by bulk solution, as suggested by Eq. S.7. The scalings between nucleation rate and protein coverage therefore depend on the time at which we evaluate them, on the amount of fibril at that time and the protein and inhibitor concentrations.
By definition, every dependent physical variable (say $\theta$, $r$ or $M$) can be expressed in terms of initial conditions and time. For example:

$$M = M(m_0, M_0, I, t), \qquad [S.8]$$

where $m_0$ and $M_0$ are the initial dissolved and insoluble protein concentration, respectively, and $I$ is the initial inhibitor concentration. Now using the property that the aggregated fibril mass is always monotonically increasing with time, we can invert equation S.8 and write time in terms of concentrations as:

$$t = t(m_0, M_0, I, M) \qquad [S.9]$$

such that every dependent variable can be expressed in terms of initial conditions and the experimentally measured $M = m_{\mathrm{tot}}\,\tilde{\mathcal{M}}$ We then have:

$$\theta = \theta(m_0, M_0, I, M) \quad \text{and} \quad r = r(m_0, M_0, I, M), \qquad [S.10]$$

and can express the scaling exponent $\sigma$ as a partial derivative at constant (zero) inhibitor concentration:

$$\sigma = \left(\frac{\partial \ln r}{\partial \ln \theta}\right)\Bigg|_{I=0,M_0,M}. \qquad [S.11]$$

Here, $m_0$, and with it $m = m_0 + M_0 - M$ is allowed to vary. On the other hand, the inhibition scaling exponent $\sigma_{\mathrm{I}}$ can be expressed as a partial derivative at constant concentration of free monomer:

$$\sigma_{\mathrm{I}} = \left(\frac{\partial \ln r}{\partial \ln \theta}\right)\Bigg|_{m_0,M_0,M}, \qquad [S.12]$$

and the inhibitor concentration is allowed to vary. Note that the concentration of free monomer is kept constant here as $m = m_0 + M_0 - M = const.$ as $m_0, M_0, M$ are all constant.

**1.4. Experimental data.** In order to evaluate experimentally the two scaling exponents for we need two separate appropriate sets of A$\beta_{42}$ aggregation experiments. To evaluate $\sigma$, the initial concentration of A$\beta_{42}$ monomer needs to be varied in absence of any inhibitor. And to evaluate $\sigma_{\mathrm{I}}$, the initial concentration of protein monomer needs to be kept constant while the concentration of Brichos inhibitor is varied.
For extracting $\sigma_{\mathrm{I}}$ we use a newly acquired dataset of A$\beta_{42}$ aggregation in the presence of proSP-C Brichos at temperature $T = 21°C$ (description of experimental procedure in section 5). Initial A$\beta_{42}$ monomer concentration was $m_0 = 3\,\mu$M, Brichos inhibitor concentrations are $I = (0, 0.125, 0.25, 0.5, 0.75, 1.0, 1.5, 2.0, 3.0)\mu$M, and all experiments were performed at estimated 1% initial fibril seed mass ($M_0 \sim 30\,$nM). We used this experimental set for two reasons. First, the initial fibril seed enables secondary nucleation to always dominate over primary nucleation, even at initial stages of the aggregation reaction, justifying our notion that the overall nucleation rate (Eq. S.3) is a good proxy for the secondary nucleation rate. And secondly, measurements of Brichos binding kinetics were performed at a similar temperature of $T = 25°C$ [5, 6].
For extracting $\sigma$ we use a previously published dataset of A$\beta_{42}$ aggregation at a temperature $T = 37°C$ published by Cohen et al. in 2013 [1]. The initial A$\beta_{42}$ monomer concentrations are $m_0 = (5.0, 4.0, 3.5, 3.0, 2.5, 2.0, 1.75, 1.5, 1.35, 1.2)\mu M$, and there is no fibril seed or any inhibitor present. We chose this dataset for its exceptional quality in terms of having no drift of ThT signal in the plateau phase of the aggregation reaction making the process of data normalisation more robust.

**1.5. Experimental analysis workflow.** The exact procedure we used to extract the scaling exponent $\sigma_{\mathrm{I}}$ at varying inhibitor is the following. First, we take our normalised aggregation dataset (datapoints in Fig. S1a)and fit the curves at different inhibitor concentrations with a closely fitting sigmoid function ($\tilde{\mathcal{M}}(I,t) = M_0/m_{\mathrm{tot}} + m_0/m_{\mathrm{tot}} * (1 + 1/(1 + a/(e^{bt} - 1)))$) (dashed lines in Fig. S1a). This allows us to take time-derivatives of $\tilde{\mathcal{M}}$ that we use to extract the rate of nucleation, and to calculate the times at which a certain amount of fibril aggregate is formed (e.g. times for different inhibitor concentrations at which $M(t) \equiv m_0 * \tilde{\mathcal{M}} = 1\mu M$, as shown in Fig. S1b). Using Eqs. S.3 and S.6-S.7 we can then directly extract the rate of nucleation (Fig. S1c) and both inhibitor and protein coverages (Fig. S1d,e). We then make a parametric plot of the nucleation rate against protein coverage on a log-log scale (Fig. S1f). And while the variables $m_0$ and $M_0$ are automatically kept constant for the entire dataset, we must further fix the amount of fibril mass, say at $M = 1\mu M$ in order to evaluate $\sigma_{\mathrm{I}}$. By doing that, we obtain ten points, one for each inhibitor concentration, on the log-log plot of rate against coverage (gray points in Fig. S1f). The inhibitor scaling exponent $\sigma_{\mathrm{I}}$ is then obtained by fitting a straight line through these points and extracting the slope, in line with the definition of Eq. S.12 (black dashed line in Fig. S1f). The workflow to extract the scaling exponent $\sigma$ is very similar and is presented in Fig. S2. To plot data in Fig. 1c of the main text, we used the values of coverage, rates, and the resulting scaling exponents at $M = 0.9\,\mu$M. Here, $\sigma = 2.07 \pm 0.04$, and $\sigma_{\mathrm{I}} = 0.98 \pm 0.02$, where the small uncertainties are due to the linear fit.

**1.6. Sensitivity analysis.** We also analyse how the scaling exponent changes with the amount of aggregated fibril mass and with the values of different rate constants. Crucially, at literature values of binding rate constants, we find that the inhibitor scaling exponent remains well-bounded around the value of $\sigma_{\rm I} = 1$ in both the nucleation and growth phases (orange data in Fig. S3a). It starts to deviate only closer to the plateau phase. The value of the scaling exponent $\sigma$ appears less stable (blue data in Fig. 3a). This is because keeping $M$ constant for a dataset with varying initial monomer concentration crosses regimes with some aggregations reactions in the nucleation/growth phase and others already in the plateau phase (see grey points in Fig. S2b). The values for the scaling exponents we quote in the main text come from averaging over $M$. The value $\sigma = 2.3 \pm 0.4$ is the mean and standard deviation of blue data in Fig. S3a. And $\sigma_{\rm I} = 1.0 \pm 0.1$ is the mean and standard deviation of orange data in Fig. S3a.

As for how much the scaling exponents are sensitive to exact values of rate constants, we first note that only the inhibitor rate constants $k_{\rm I}^{+}$ and $k_{\rm I}^{-}$ play a significant role. The elongation rate constant $k_{+}$ appears as an additive constant under the logarithm so it does not influence the values of the scaling exponents. And the same is true for the protein dissociation constant $K_{\rm d}$ at dilute conditions, where $m \ll K_{\rm d}$ implies $\theta_0 \approx m/K_{\rm d}$ and then $K_{\rm d}$ also only appear only as an additive constant. In Fig. S3b, we plot how the extracted inhibition scaling exponent $\sigma_{\rm I}$ depends on the value of inhibitor dissociation constant $K_{\rm I} = k_{\rm I}^{-}/k_{\rm I}^{+}$ where both on and off rate constant are allowed to vary at the same time. We find that even under high uncertainty in $K_{\rm I} = 130\,\text{nm} \pm 50\%$, the value of $\sigma_{\rm I}$ remains close to unity.

### 2. Statistical mechanics models of secondary nucleation and its inhibition

In this section we develop an equilibrium statistical mechanics model of secondary nucleation where two species (a nucleating species and inhibitor) are competing for the same surface. We first explore the general case and derive the conditions under which secondary nucleation rate scales linearly with protein coverage when varying inhibitor concentration ($\sigma_\mathrm{I} = 1$). We then solve the model for the simple geometry of the catalytic surface where catalytic sites are very confined, able to accommodate only up to two adsorbing particles. Next, we treat a general class of models that observe the requirement $\sigma_\mathrm{I} = 1$ and derive a formula for competitive binding. We then use this formula to express equilibrium stat-mech models of secondary rate in a form that can be translated into mass-action kinetic rate laws. Finally, we discuss alternative scenarios under which $\sigma_\mathrm{I} = 1$ can be satisfied.

**2.1. General model.** We treat secondary nucleation as a two-step process involving 1) the formation of a pre-nucleating cluster (protein oligomer) on the fibril surface, and 2) its subsequent catalysis into a fibril nucleus [7–9]. Our model does not presume where the formation of pre-nucleating clusters takes place - it can be in the bulk solution or directly on the fibril surface. We only need to recognise that the catalytic step takes place in the presence of the surface and assume that catalysis happens on a slower timescale than monomer rearrangement into clusters.

As described in the main text, we define all separate areas of fibril surface where monomers and inhibitors can bind as an ensemble of independent subsystems. The subsystem A can be considered independent from subsystem B if there are no correlations in adsorbent occupancies between A and B. This if surely true if the subsystems A and B correspond to two distinct amyloid fibrils. But it can also be true if the subsystems A and B are distinct catalytic sites that are part of the same fibril. In this case, A and B can be considered independent under the condition that they are far apart from each other such that there are no interactions between adsorbents bound to different sites. But also the movement of the fibril needs to be slow compared to the adsorption/desorption timescales such that the system can be considered to be at equilibrium.

Every catalytic area is treated as equivalent and is able to accommodate up to $N_\mathrm{max}$ particles such that any number of nucleating monomers ($N_\mathrm{m}$) and inhibitors ($N_\mathrm{I}$) can adsorb to the catalytic site subject to a constraint $N_\mathrm{m} + N_\mathrm{I} \leq N_\mathrm{max}$. The adsorption of both species and their clustering on a given catalytic site is governed by a per-area grand-canonical partition function $\xi(\lambda_\mathrm{m}, \lambda_\mathrm{I}, N_\mathrm{max}, T)$, where $\lambda_\mathrm{m}$ and $\lambda_\mathrm{I}$ are the monomer and inhibitor thermodynamic activities, respectively, and $T$ is the temperature. Generally, we have:

$$\xi = \sum_{N_\mathrm{m}=0}^{N_\mathrm{max}} \sum_{N_\mathrm{I}=0}^{N_\mathrm{max}-N_\mathrm{m}} Q_{N_\mathrm{m},N_\mathrm{I}} \lambda_\mathrm{m}^{N_\mathrm{m}} \lambda_\mathrm{I}^{N_\mathrm{I}}, \tag{S.13}$$

where $Q_{N_\mathrm{m},N_\mathrm{I}}$ is the partition function for the case with $N_\mathrm{m}$ monomers and $N_\mathrm{I}$ inhibitors bound to the catalytic area. The coverage of the catalytic surface by the nucleating species ($\theta \equiv \langle N_\mathrm{m} \rangle / L$) can be extracted from the grand partition function as:

$$\theta = \frac{1}{N_\mathrm{max}\, \xi} \sum_{N_\mathrm{m}=0}^{N_\mathrm{max}} \sum_{N_\mathrm{I}=0}^{N_\mathrm{max}-N_\mathrm{m}} N_\mathrm{m}\, Q_{N,N_\mathrm{I}} \lambda_\mathrm{m}^{N_\mathrm{m}} \lambda_\mathrm{I}^{N_\mathrm{I}}, \tag{S.14}$$

and the rate of secondary nucleation due to a single catalytic area can be similarly expressed as:

$$r = \frac{1}{\xi} \sum_{N_\mathrm{m}=0}^{N_\mathrm{max}} \sum_{N_\mathrm{I}=0}^{N_\mathrm{max}-N_\mathrm{m}} r_{N_\mathrm{m},N_\mathrm{I}}\, Q_{N_\mathrm{m},N_\mathrm{I}} \lambda_\mathrm{m}^{N_\mathrm{m}} \lambda_\mathrm{I}^{N_\mathrm{I}}, \tag{S.15}$$

where $r_{N_\mathrm{m},N_\mathrm{I}}$ is the average rate at a given occupancy of monomers and inhibitors. This per-occupancy rate is in turn given by a canonical ensemble average over all different configurations $i$ that $N_\mathrm{m}$ monomers and $N_\mathrm{I}$ inhibitors can arrange into:

$$r_{N_\mathrm{m},N_\mathrm{I}} = \frac{1}{Q_{N_\mathrm{m},N_\mathrm{I}}} \sum_i e^{-E_i/kT} \sum_{j=2}^{N_\mathrm{m}} n_i(j) \mathcal{C}_i(j). \tag{S.16}$$

Here, $E_i$ is the energy of a given configuration, $n_i(j)$ is the protein cluster distribution at configuration $i$ (with $j$ the size of the cluster), and $\mathcal{C}_i(j)$ is the rate of catalysis for a protein cluster of size $j$ at the configuration $i$. This form of the rate accounts for the possibility that protein oligomers of different sizes and shapes get catalysed at different rates, and a further possibility that the presence of inhibitors affect the catalysis rate. Combining the formulas for the average rate and coverage (S.14-S.16), we can evaluate the scaling exponents $\sigma$ and $\sigma_\mathrm{I}$ by keeping $\lambda_\mathrm{I} = 0$ or $\lambda_\mathrm{m}$ at a constant value, respectively.

Regardless of the potentially complicated details of the nucleating system, as captured in the general form of $Q_{N_\mathrm{m},N_\mathrm{I}}$ and $\mathcal{C}_i(j)$, we can find a condition under which $\sigma_\mathrm{I} = 1$. To do this, we separate the grand-canonical sum over all possible occupancies $N_\mathrm{m}$ and $N_\mathrm{I}$ into four parts:

$$\sum_{N_\mathrm{m}=0}^{N_\mathrm{max}} \sum_{N_\mathrm{I}=0}^{N_\mathrm{max}-N_\mathrm{m}} \equiv 1 + \sum_{N_\mathrm{m}=1}^{N_\mathrm{max}} + \sum_{N_\mathrm{I}=1}^{N_\mathrm{max}} + \sum_{N_\mathrm{m}=1}^{N_\mathrm{max}} \sum_{N_\mathrm{I}=1}^{N_\mathrm{max}-N_\mathrm{m}} . \tag{S.17}$$

Here the first term describes a catalytic area that is devoid of any adsorbents. The second and third terms describe solo occupancies of the catalytic area by either protein monomers or inhibitors, respectively. And the last term describes situations where both monomers and inhibitors are co-occupying the catalytic area. In this way, we can express the protein coverage as:

$$\theta = \frac{1}{N_{\max}\,\xi}\left\{ \sum_{N_\mathrm{m}=1}^{N_{\max}} N_\mathrm{m}\, Q_{N_\mathrm{m},0}\lambda_\mathrm{m}^{N_\mathrm{m}} + \sum_{N_\mathrm{m}=1}^{N_{\max}} \sum_{N_\mathrm{I}=1}^{N_{\max}-N_\mathrm{m}} N_\mathrm{m}\, Q_{N_\mathrm{m},N_\mathrm{I}}\lambda_\mathrm{m}^{N_\mathrm{m}}\lambda_\mathrm{I}^{N_\mathrm{I}} \right\}, \quad \text{[S.18]}$$

and the rate as:

$$r = \frac{1}{\xi}\left\{ \sum_{N_\mathrm{m}=1}^{N_{\max}} r_{N_\mathrm{m},0}\, Q_{N_\mathrm{m},0}\lambda_\mathrm{m}^{N_\mathrm{m}} + \sum_{N_\mathrm{m}=1}^{N_{\max}} \sum_{N_\mathrm{I}=1}^{N_{\max}-N_\mathrm{m}} r_{N_\mathrm{m},N_\mathrm{I}}\, Q_{N_\mathrm{m},N_\mathrm{I}}\lambda_\mathrm{m}^{N_\mathrm{m}}\lambda_\mathrm{I}^{N_\mathrm{I}} \right\}. \quad \text{[S.19]}$$

Putting both the rate and coverage into an expression for $\sigma_\mathrm{I}$ gives:

$$\sigma_\mathrm{I} \equiv \left(\frac{\partial \ln r}{\partial \ln \theta}\right)_{\lambda_\mathrm{m},N_{\max},T} = \left(\frac{\partial \ln\left[\sum_{N_\mathrm{m}=1}^{N_{\max}} r_{N_\mathrm{m},0}\, Q_{N_\mathrm{m},0}\lambda_\mathrm{m}^{N_\mathrm{m}} + \sum_{N_\mathrm{m}=1}^{N_{\max}}\sum_{N_\mathrm{I}=1}^{N_{\max}-N_\mathrm{m}} r_{N_\mathrm{m},N_\mathrm{I}}\, Q_{N_\mathrm{m},N_\mathrm{I}}\lambda_\mathrm{m}^{N_\mathrm{m}}\lambda_\mathrm{I}^{N_\mathrm{I}}\right] - \partial\ln\xi}{\partial \ln\left[\sum_{N_\mathrm{m}=1}^{N_{\max}} N_\mathrm{m}\, Q_{N_\mathrm{m},0}\lambda_\mathrm{m}^{N_\mathrm{m}} + \sum_{N_\mathrm{m}=1}^{N_{\max}}\sum_{N_\mathrm{I}=1}^{N_{\max}-N_\mathrm{m}} N_\mathrm{m}\, Q_{N_\mathrm{m},N_\mathrm{I}}\lambda_\mathrm{m}^{N_\mathrm{m}}\lambda_\mathrm{I}^{N_\mathrm{I}}\right] - \partial\ln\xi}\right)_{\lambda_\mathrm{m},N_{\max},T}. \quad \text{[S.20]}$$

In both the numerator and denominator, the first term is made out of two sums while the second term is the same ($\partial \ln \xi$). The first sum of the first term (over solo protein occupancies) evidently depends only on the monomer activity and is kept constant, while the double-sum (over co-occupancies) depend both on monomer and inhibitor activities. So in order to have the first terms of the numerator and denominator disappear we require that both double-sums over co-occupancies are equal to zero. In that case we have

$$\sigma_\mathrm{I} = \left(\frac{\partial f(\lambda_\mathrm{m}) - \partial \ln \xi(\lambda_\mathrm{m},\lambda_\mathrm{I})}{\partial g(\lambda_\mathrm{m}) - \partial \ln \xi(\lambda_\mathrm{m},\lambda_\mathrm{I})}\right)_{\lambda_\mathrm{m}} = 1, \quad \text{[S.21]}$$

where $f$ and $g$ denote the first sums of the numerator and denominator, respectively. Notably, the double-sums vanish under a common condition: $Q_{N_\mathrm{m}\geq 1, N_\mathrm{I}\geq 1} = 0$. So a linear scaling $\sigma_\mathrm{I}$ emerges only if there is no co-occupation of monomers and inhibitors on the same catalytic area. This can for example be achieved if all configurations with monomer-inhibitor co-occupancy are prohibited by large energies $E_i >> 0$. In the context of a fibril surface and considering only short-range interactions between adsorbates such large positive co-occupation energies would likely only originate from steric interactions. This leads us to conclude that a measured scaling exponent of $\sigma_\mathrm{I} = 1$ is compatible with a very confined geometry of catalytic areas where co-occupancy is restricted by hard-core repulsion (volume exclusion) between monomers and inhibitors. We call such confined catalytic areas "catalytic sites" which have the properties of being small (no co-occupancy) and distant (they are independent subsystems).

**2.2. Small and distant catalytic sites model.** Knowing the conditions for which we get $\sigma_\mathrm{I} = 1$ we can now build a simple equilibrium model of secondary nucleation and its inhibition on confined catalytic sites. To do this, we consider the smallest catalytic site of size $N_{\max} = 2$ that can theoretically still support nucleation through a pre-nucleation cluster. According to Eq. S.13 the per-site grand partition function for this simple system reads:

$$\xi^{N_{\max}=2} = 1 + Q_{1,0}\lambda_\mathrm{m} + Q_{0,1}\lambda_\mathrm{I} + Q_{2,0}\lambda_\mathrm{m}^2 + Q_{0,2}\lambda_\mathrm{I}^2 + Q_{1,1}\lambda_\mathrm{m}\lambda_\mathrm{I}, \quad \text{[S.22]}$$

which can be evaluated as:

$$\xi^{N_{\max}=2} = 1 + 2q_\mathrm{m}\lambda_\mathrm{m} + 2q_\mathrm{I}\lambda_\mathrm{I} + q_\mathrm{m}^2\lambda_\mathrm{m}^2 + q_\mathrm{I}^2\lambda_\mathrm{I}^2 + 2q_\mathrm{m}q_\mathrm{I}\lambda_\mathrm{m}\lambda_\mathrm{I}\, e^{-\epsilon/kT}. \quad \text{[S.23]}$$

Here, $q_\mathrm{m}$ and $q_\mathrm{I}$ are the molecular partition functions for the sole adsorbed protein monomer and inhibitor, respectively, with $\epsilon_\mathrm{mI}$ the interaction energy between monomer and inhibitor or when they are co-occupying the catalytic site. We neglect monomer-monomer and inhibitor-inhibitor interactions. For $N_{\max} = 2$ the only pre-nucleation cluster that can be catalysed is a protein dimer and the rate of nucleation is now given simply by:

$$r^{N_{\max}=2} = \frac{\mathcal{C}\, q_\mathrm{m}^2\lambda_\mathrm{m}^2}{\xi^{N_{\max}=2}}, \quad \text{[S.24]}$$

where $\mathcal{C}$ is a catalysis rate constant. The protein coverage can be evaluated as:

$$\theta^{N_{\max}=2} = \frac{q_\mathrm{m}\lambda_\mathrm{m} + q_\mathrm{m}q_\mathrm{I}\lambda_\mathrm{m}\lambda_\mathrm{I}\, e^{-\epsilon/kT} + q_\mathrm{m}^2\lambda_\mathrm{m}^2}{\xi^{N_{\max}=2}}. \quad \text{[S.25]}$$

In the absence of inhibitor, the scaling exponent at varying $\lambda_\mathrm{m}$ is:

$$\sigma^{N_{\max}=2} = \left(\frac{\partial \ln r^{N_{\max}=2}}{\partial \ln \theta^{N_{\max}=2}}\right)_{\lambda_\mathrm{I}=0,N_{\max},T} = \left(\frac{\partial \ln(q_\mathrm{m}\lambda_\mathrm{m}/(1+q_\mathrm{m}\lambda_\mathrm{m}))^2}{\partial \ln(q_\mathrm{m}\lambda_\mathrm{m}/(1+q_\mathrm{m}\lambda_\mathrm{m}))}\right)_{\lambda_\mathrm{I}=0,N_{\max},T} = 2. \quad \text{[S.26]}$$

And the inhibition scaling exponent as:

$$\sigma_{\mathrm{I}}^{N_{\max}=2} = \left(\frac{\partial \ln r^{N_{\max}=2}}{\partial \ln \theta^{N_{\max}=2}}\right)_{\lambda_{\mathrm{m}}, N_{\max}, T} = \left(\frac{\partial \ln(q_{\mathrm{m}}^2 \lambda_{\mathrm{m}}^2) - \partial \ln(\xi^{N_{\max}=2})}{\partial \ln(q_{\mathrm{m}}\lambda_{\mathrm{m}} + q_{\mathrm{m}}^2\lambda_{\mathrm{m}}^2 + q_{\mathrm{m}} q_{\mathrm{I}} \lambda_{\mathrm{m}} \lambda_{\mathrm{I}}\, e^{-\epsilon/kT}) - \partial \ln(\xi^{N_{\max}=2})}\right)_{\lambda_{\mathrm{m}}, N_{\max}, T}. \quad [S.27]$$

As presented in the main text (Fig. 3b), the value of $\sigma_{\mathrm{I}}$ depends on the interaction energy between the co-occupying monomer and inhibitor. At $\epsilon = 0$ we find $\sigma_{\mathrm{I}}^{N_{\max}=2} = \sigma^{N_{\max}=2} = 2$, and for infinitely repulsive interaction $\epsilon \to \infty$, we have $\sigma_{\mathrm{I}}^{N_{\max}=2} = 1$, consistent with the analysis of section 2.1. At intermediate repulsion $\epsilon > 0$ we find $1 < \sigma_{\mathrm{I}}^{N_{\max}=2} < \sigma^{N_{\max}=2}$, and for attractive interaction $\epsilon < 0$ we have $\sigma_{\mathrm{I}}^{N_{\max}=2} > \sigma^{N_{\max}=2}$. The pink solid line in Fig. 3b of the main text is the analytical expression given by Eq. S.27. The green line is likewise an analytical expression but for $N_{\max} = 3$. We have also analytically derived $\sigma_{\mathrm{I}}$ for $N_{\max} = 4$ and confirmed the expression is almost indistinguishable from that of $N_{\max} = 3$ (not shown).

**2.3. Inhibitor covers entire catalytic site model.** We now turn to a class of small catalytic sites models that automatically result in $\sigma_{\mathrm{I}} = 1$ and that can serve as a basis for mass-action formulas that we use to model the secondary nucleation kinetics of $A\beta_{42}$ in the presence or absence of Brichos. Here, we already take into account that monomers and inhibitors cannot co-occupy a catalytic site, and additionally model the inhibitor as covering the whole catalytic site on its own (which is anyway somewhat more realistic than expecting that e.g. a monomer and inhibitor cannot co-occupy a site while two inhibitors can). We have for the per-site grand-partition function:

$$\xi^C = 1 + Q_{\mathrm{I}}\lambda_{\mathrm{I}} + \sum_{N_{\mathrm{m}}=1}^{N_{\max}} Q_{N_{\mathrm{m}}} \lambda_{\mathrm{m}}^{N_{\mathrm{m}}} \quad [S.28]$$

For the coverages we then have:

$$\theta_{\mathrm{I}} = \frac{Q_{\mathrm{I}}\lambda_{\mathrm{I}}}{\xi^C}, \qquad \theta = \frac{1}{N_{\max}\xi^C} \sum_{N_{\mathrm{m}}=1}^{N_{\max}} N_{\mathrm{m}} Q_{N_{\mathrm{m}}} \lambda_{\mathrm{m}}^{N_{\mathrm{m}}}, \quad [S.29]$$

where $\theta_{\mathrm{I}}$ is the inhibitor coverage, and $\theta$ the protein coverage. The rate of nucleation here is:

$$r = \frac{1}{\xi^C} \sum_{N_{\mathrm{m}}=2}^{N_{\max}} r_{N_{\mathrm{m}}} Q_{N_{\mathrm{m}}} \lambda_{\mathrm{m}}^{N_{\mathrm{m}}}, \quad [S.30]$$

which can be expressed, using Eq. S.29, as:

$$r = \theta \cdot f(\theta_0). \quad [S.31]$$

Here $f(\theta_0)$ denotes that $f$ is a function of the protein coverage in absence of inhibitor, and we have taken into account that $\theta_0$ is a monotonically increasing function of $\lambda_{\mathrm{m}}$.

**2.4. Competitive binding.** Taking together Eqs. S.28 and S.29, we can derive that the protein coverage can be factorised as:

$$\theta(\lambda_{\mathrm{m}}, \lambda_{\mathrm{I}}) = (1 - \theta_{\mathrm{I}}(\lambda_{\mathrm{m}}, \lambda_{\mathrm{I}}) \cdot \theta_0(\lambda_{\mathrm{m}}). \quad [S.32]$$

Here, $\theta_0 \equiv \theta(I = 0)$ is again the protein coverage in absence of any inhibitor, and $(1 - \theta_{\mathrm{I}})$ is the proportional amount of surface not occupied by inhibitor. This expression for competitive binding (Eq. S.32) also holds exactly the same in the case of ideal monomer-inhibitor interactions. This result is crucial as it suggests that the effect of competitive binding can be expressed in the same manner regardless of the exact inhibition mechanism. Moreover, in the case of either low protein coverage $\theta_0 \ll 1$ or when $\theta_{\mathrm{I}} \gg \theta_0$, overall protein coverage can be additionally simplified as

$$\theta(\lambda_{\mathrm{m}}, \lambda_{\mathrm{I}}) \approx (1 - \theta_{\mathrm{I}}(\lambda_{\mathrm{I}})) \cdot \theta_0(\lambda_{\mathrm{m}}), \quad [S.33]$$

where $\theta_{\mathrm{I}}$ is now only a function of inhibitor activity. This expression is very useful because it tells us that the protein coverage can be evaluated by knowing the individual binding behaviour of monomers and inhibitors separately from one another. We use this when experimentally extracting the inhibition scaling exponent from the aggregation kinetics of $A\beta_{42}$ in the presence of Brichos where the binding kinetics of the protein and inhibitor are known only in isolation.

**2.5. Application to chemical kinetics of A$\beta$ secondary nucleation.** We now use the results from the above subsections to derive rate formulas that can be used for modelling the kinetics of secondary nucleation and its inhibition.
First, we use the expression for competitive binding (Eq. S.32), and express the rate for the small catalytic sites model as

$$r_{\text{small sites}} = (1 - \theta_{\mathrm{I}})\, g(\theta_0), \quad [S.34]$$

where $g(\theta_0) = \theta_0 f(\theta_0)$ is a function that depends on the specific secondary nucleation mechanism. We therefore find that in the case of inhibitors covering whole catalytic sites, the inhibition mechanism is expressed as a simple linear factor $(1 - \theta_{\mathrm{I}})$ that represents the proportion of catalytic sites that are available for secondary nucleation. This is a general result, independent of the form of the detailed expressions for $r_{N_{\mathrm{m}}}$, and $Q_{N_{\mathrm{m}}}$.

In contrast to the above result, in the case of the ideal Langmuir inhibition mechanism, the inhibitor acts by renormalising the amount of protein bound to a given catalytic site (instead of taking some catalytic sites away) according to Eq. S.32, and the rate can be expressed as:

$$r_{\mathrm{ideal}} = g(\theta). \tag{S.35}$$

So we see, that the rate of secondary nucleation can be expressed in the absence of inhibitor as $r = g(\theta_0)$. Previous work has shown that in the case of A$\beta$ aggregation, the function $g$ simply takes the form $g(\theta_0) = \theta_0^{n_2}$, where $n_2$ is the reaction order [4, 9, 10]. However, taking into account the different inhibition mechanism, we have $r_{\text{small sites}} = (1-\theta_{\mathrm{I}})\,\theta_0^{\sigma}$, and $r_{\mathrm{ideal}} = ((1-\theta_{\mathrm{I}})\,\theta_0)^{\sigma}$, where we have made a substitution $n_2 = \sigma$, which is true by definition. We see that the difference between the above two rate equations is only the value of the scaling exponent of the factor $(1-\theta_{\mathrm{I}})$. We can therefore express the rate for both inhibition mechanisms with a single formula as:

$$r = (1-\theta_{\mathrm{I}})^{\sigma_{\mathrm{I}}}\,\theta_0^{\sigma}, \tag{S.36}$$

where for the small catalytic sites model $\sigma_{\mathrm{I}} = 1$ and for ideal binding model $\sigma_{\mathrm{I}} = \sigma$.

**2.6. Alternative nucleation scenarios.** Here we present and discuss mechanistic scenarios that might lead to the behaviour of the inhibition scaling exponent as $\sigma_{\mathrm{I}} = 1$.

**Stacking model:** First we discuss the possibility that protein oligomers form by monomer stacking on top of one another (see Fig. S5 for schematic). The reasoning for why we could get $\sigma_{\mathrm{I}} = 1$ here is that a stacked oligomer might form as a simple reaction between a surface-bound A$\beta$ monomer and a monomer from solution. This would give a rate formula of the form $r \propto \theta\, f(\lambda_{\mathrm{m}})$, where is A$\beta$ coverage is dependent on both monomer ($\lambda_{\mathrm{m}}$) and inhibitor ($\lambda_{\mathrm{I}}$) concentration (or thermodynamic activity to be more exact) while the function $f(\lambda_{\mathrm{m}})$ is only dependent on free monomer in solution. And one might think that this rate formula would be true for stacking regardless of the shape of the fibril surface.
We test this intuitive prediction by building on our statistical mechanics framework for describing secondary nucleation and its inhibition. Here we extend the general model of Eq. S.13 as:

$$\xi = \sum_{N_{\mathrm{m}}=0}^{N_{\max}} \sum_{N_{\mathrm{d}}=0}^{N_{\mathrm{m}}} \sum_{N_{\mathrm{I}}=0}^{N_{\max}-N_{\mathrm{m}}} Q_{N_{\mathrm{m}},N_{\mathrm{d}},N_{\mathrm{I}}} \lambda_{\mathrm{m}}^{N_{\mathrm{m}}+N_{\mathrm{d}}} \lambda_{\mathrm{I}}^{N_{\mathrm{I}}}, \tag{S.37}$$

where $N_{\mathrm{d}} \leq N_{\mathrm{m}}$ is the amount of monomers that are stacked upon existing surface-bound monomers. Here we limit ourselves to only dimers but the scheme can be easily extended to higher-order stacked oligomers. We model the rate of nucleation as proportional to the abundance of stacked dimers:

$$r = \frac{1}{\xi} \sum_{N_{\mathrm{m}}=0}^{N_{\max}} \sum_{N_{\mathrm{d}}=0}^{N_{\mathrm{m}}} \sum_{N_{\mathrm{I}}=0}^{N_{\max}-N_{\mathrm{m}}} N_{\mathrm{d}}\, Q_{N_{\mathrm{m}},N_{\mathrm{d}},N_{\mathrm{I}}} \lambda_{\mathrm{m}}^{N_{\mathrm{m}}+N_{\mathrm{d}}} \lambda_{\mathrm{I}}^{N_{\mathrm{I}}}, \tag{S.38}$$

and the total coverage of the surface by protein as:

$$\theta = \frac{1}{2N_{\max}\xi} \sum_{N_{\mathrm{m}}=0}^{N_{\max}} \sum_{N_{\mathrm{d}}=0}^{N_{\mathrm{m}}} \sum_{N_{\mathrm{I}}=0}^{N_{\max}-N_{\mathrm{m}}} (N_{\mathrm{m}}+N_{\mathrm{d}})\, Q_{N_{\mathrm{m}},N_{\mathrm{d}},N_{\mathrm{I}}} \lambda_{\mathrm{m}}^{N_{\mathrm{m}}+N_{\mathrm{d}}} \lambda_{\mathrm{I}}^{N_{\mathrm{I}}}. \tag{S.39}$$

As before, we can separate the grand-canonical sums into multiple parts:

$$\sum_{N_{\mathrm{m}}=0}^{N_{\max}} \sum_{N_{\mathrm{d}}=0}^{N_{\mathrm{m}}} \sum_{N_{\mathrm{I}}=0}^{N_{\max}-N_{\mathrm{m}}} \equiv 1 + \sum_{N_{\mathrm{m}}=1}^{N_{\max}} \sum_{N_{\mathrm{d}}=0}^{N_{\mathrm{m}}} + \sum_{N_{\mathrm{I}}=1}^{N_{\max}} + \sum_{N_{\mathrm{m}}=1}^{N_{\max}} \sum_{N_{\mathrm{d}}=0}^{N_{\mathrm{m}}} \sum_{N_{\mathrm{I}}=1}^{N_{\max}-N_{\mathrm{m}}}. \tag{S.40}$$

Putting both the rate and coverage into an expression for $\sigma_{\mathrm{I}}$ gives

$$\sigma_{\mathrm{I}} = \left( \frac{\partial \ln \left[ \sum_{N_{\mathrm{m}}=1}^{N_{\max}} \sum_{N_{\mathrm{d}}=1}^{N_{\mathrm{m}}} N_{\mathrm{d}}\, Q_{N_{\mathrm{m}},N_{\mathrm{d}}} \lambda_{\mathrm{m}}^{N_{\mathrm{m}}+N_{\mathrm{d}}} + \sum_{N_{\mathrm{m}}=1}^{N_{\max}} \sum_{N_{\mathrm{d}}=1}^{N_{\mathrm{m}}} \sum_{N_{\mathrm{I}}=1}^{N_{\max}-N_{\mathrm{m}}} N_{\mathrm{d}}\, Q_{N_{\mathrm{m}},N_{\mathrm{d}},N_{\mathrm{I}}} \lambda_{\mathrm{m}}^{N_{\mathrm{m}}+N_{\mathrm{d}}} \lambda_{\mathrm{I}}^{N_{\mathrm{I}}} \right] - \partial \ln \xi}{\partial \ln \left[ \sum_{N_{\mathrm{m}}=1}^{N_{\max}} \sum_{N_{\mathrm{d}}=0}^{N_{\mathrm{m}}} (N_{\mathrm{m}}+N_{\mathrm{d}})\, Q_{N_{\mathrm{m}},N_{\mathrm{d}}} \lambda_{\mathrm{m}}^{N_{\mathrm{m}}+N_{\mathrm{d}}} + \sum_{N_{\mathrm{m}}=1}^{N_{\max}} \sum_{N_{\mathrm{d}}=0}^{N_{\mathrm{m}}} \sum_{N_{\mathrm{I}}=1}^{N_{\max}-N_{\mathrm{m}}} (N_{\mathrm{m}}+N_{\mathrm{d}})\, Q_{N_{\mathrm{m}},N_{\mathrm{d}},N_{\mathrm{I}}} \lambda_{\mathrm{m}}^{N_{\mathrm{m}}+N_{\mathrm{d}}} \lambda_{\mathrm{I}}^{N_{\mathrm{I}}} \right] - \partial \ln \xi} \right)_{\lambda_{\mathrm{m}}}. \tag{S.41}$$

Remarkably, we find that the condition under which $\sigma_{\mathrm{I}} = 1$ is the same no co-occupancy condition as for the model without stacking:

$$\sigma_{\mathrm{I}} = 1 \qquad \textbf{if} \qquad Q_{N_{\mathrm{m}}\geq 1, N_{\mathrm{d}}, N_{\mathrm{I}}\geq 1} = 0. \tag{S.42}$$

This means we should not expect to have $\sigma_{\mathrm{I}} = 1$ in case a) of Fig. S4, where the whole fibril surface is a single large catalytic area and inhibitors can obviously co-occupy it with monomers. Also in the case b) of Fig. S5, where binding sites are localised instead of diffuse, the no co-occupancy criterion can be satisfied only if the binding sites can be considered independent. That would be the case only in the very ideal scenario where bound inhibitors and monomers would not interact in any way between the adjacent binding sites. Which is only the implausible scenario where monomers and inhibitors would be of the exact same size as the localised binding site and have no additional interactions with their neighbours. Finally, in c) of Fig.

S5, the no co-occupancy criterion is naturally observed and the scenario reduces to nucleation on small and distant catalytic sites.

**Monomer catalysis** Another alternative model consistent with $\sigma_{\mathrm{I}} = 1$ would be a secondary nucleation mechanism that does not involve the presence of oligomers on the fibril surface. Instead, it could involve only binding of protein monomers to the fibril surface which would get catalysed in their monomeric form (as opposed to an oligomer). Such a 'catalysed' monomer might then detach, and grow further in solution until a fibril nucleus is formed. In that case the secondary nucleation rate formula would again be $r \propto \theta\, f(\lambda_{\mathrm{m}})$, regardless of the properties of the surface. However, previous studies showed that on-pathway oligomers that will eventually turn into daughter fibrils can indeed be seen under the microscope to form at the surface of a parent fibril [11, 12].

**2.7. Further applications.** In this work, we focused mostly on the monomer-dependent secondary nucleation where the catalyst and the substrate are of the same protein species. This process is distinct from heterogeneous primary nucleation only in that no foreign surface is involved in the catalytic process. However, our equilibrium statistical-mechanical model does not reference the origin of the catalytic surface, only its necessary presence. So our model should be applicable to all types of surface-based two-step nucleation processes that involve the nucleation or catalysis of species A in the presence of a non-nucleating competing species B.

### 3. Computer model

In our coarse-grained computer model of secondary nucleation, each particle is represented by a spherocylinder with a length to width ratio of $l/w = 3$ which mimics the elementary $\beta$-sheet unit in A$\beta$ peptides with dimensions 6 nm × 2 nm. The unit $w = 2$ nm defines the natural distance measure in our model against which every other distance is compared. A hard core repulsion between all particles forbids any distance between any two points on spherocylinder centerlines to be smaller than $w$.

**3.1. Interactions.** The attractive potential $V_{\text{ss}}$ between two monomers in the dissolved '$s$'-state is implemented as:

$$V_{\text{ss}}(r) = \begin{cases} -\epsilon_{\text{ss}} \left(\frac{w}{r}\right)^6 & \text{if } r \leq r_{\text{cut}} \\ 0 & \text{if } r > r_{\text{cut}}, \end{cases} \qquad \text{[S.43]}$$

where $r$ is the distance between the centers of the attractive tips located at the spherocylinder's ends, $r_{\text{cut}} = 1.3w$ is the cutoff distance, and $\epsilon_{\text{ss}}$ is the minimum interaction energy between two dissolved monomers. An attractive patch is added to only one of the two spherocylinder poles to ensure formation of finite micellar-like oligomers where the tips of dissolved monomers are all in contact at the oligomer center (see Fig. 2). The tip-to-tip interaction between any two tipped monomers is implemented in the same way (Eq. S.43) but with different interaction strengths: $\epsilon_{\text{ss}} \to \epsilon_{\text{ii}}$ for the interactions between two intermediate monomers, and $\epsilon_{\text{ss}} \to \epsilon_{\text{si}}$ for the interaction between a '$s$'-state monomer and an '$i$'-state monomer.
The attractive side-patch of the $\beta$-state monomer is $l_{\text{b}} = 0.7l$ long and spans an angle of 180°. If patches of two $\beta$-state monomers face each other their interaction $V_{\beta\beta}$ is:

$$V_{\beta\beta}(r) = \begin{cases} -\epsilon_{\beta\beta}\, cos^2(\phi) - \epsilon_{\beta\beta} \left(\frac{w}{r}\right) & \text{if } d \leq r_{\text{cut}} \\ 0 & \text{if } d > r_{\text{cut}}, \end{cases} \qquad \text{[S.44]}$$

where $2\epsilon_{\beta\beta}$ is the maximal interaction strength between two $\beta$-particles, $\phi$ is the angle between the long axes of the particles, $d$ is the shortest distance between the axes of the patches, and $r$ is the distance between the centers of the patches. The first term ensures that proteins in the $\beta$-form pack parallel to each other, mimicking the hydrogen-bond interactions between $\beta$-sheets, while the second term promotes compactness of the fibrils. This is also the interaction that holds together the mature fibril at the centre of our simulation box. To make fibrils thermodynamically stable, $V_{\beta\beta}$ has to be by far the strongest of all the interactions in the system.

The cross-interaction $V_{\text{s}\beta}$ between the dissolved and the $\beta$-sheet configuration is implemented as a potential well:

$$V_{\text{s}\beta}(d) = \begin{cases} -\epsilon_{\text{s}\beta} & \text{if } d < r_{\text{cut}} \\ 0 & \text{if } d > r_{\text{cut}}, \end{cases} \qquad \text{[S.45]}$$

where $d$ is the shortest distance between the centre of the attractive tip and the axis of the $\beta$-patch, and $\epsilon_{\text{s}\beta}$ is the interaction strength. The dissolved tip has to face the 180° opening of the $\beta$-particle side patch. The $i$-$\beta$ interaction is described in the same way, with $\epsilon_{\text{s}\beta} \to \epsilon_{\text{i}\beta}$. The interactions described above (Eqs. S.43, S.44, S.45) are all required to have spontaneous nucleation at physiological conditions.
For secondary nucleation and its inhibition we additionally need binding to the fibril surface. Adsorption of the dissolved protein onto the preformed fibril is given by:

$$V_{\text{sf}}(d) = \begin{cases} -\epsilon_{\text{sf}} \left(\frac{w}{d}\right)^6 & \text{if } r \leq r_{\text{cut}} \\ 0 & \text{if } r > r_{\text{cut}} \end{cases} \qquad \text{[S.46]}$$

where $d$ is the shortest distance between the centre of the attractive tip on the dissolved protein and the fibril particle's adsorption patch with length $l_a$. Intermediate '$i$' protein and inhibitor adsorptions onto the fibril are described in the same way (Eq. S.46), with $\epsilon_{\text{sf}} \to \epsilon_{\text{if}}$, and $\epsilon_{\text{sf}} \to \epsilon_{\text{If}}$, respectively.

**3.2. Choice of interaction parameters.** As our model contains many different interactions and particle species it can exhibit different phenomenologies depending on specific values of interaction parameters. In our simulations, we are targeting a regime where the process of self-replication dominates over spontaneous nucleation in solution.
We set the internal free energy of the $\beta$-state to 20 kT, the internal free energy of the intermediate state to 10 kT, and of the dissolved state to zero. The free energy penalty of conversion from a dissolved to $\beta$-particle is therefore $\Delta\mu_{\text{s}\to\beta} = 20$ kT while the conversion penalties to and from the intermediate state are $\Delta\mu_{\text{s}\to\text{i}} = \Delta\mu_{\text{i}\to\beta} = 10$ kT. These numbers follow the fact that aggregation-prone peptides such as A$\beta$ are typically not found in the $\beta$-sheet configuration in solution.
We choose a relatively low value of '$s$'-state binding interaction $\epsilon_{\text{ss}} = 4$ kT even though the nucleation rate is faster at larger $\epsilon_{\text{ss}}$ where oligomers would be larger and more long-lived. This was done in order to hinder spontaneous nucleation in solution whilst still retaining high rates of surface-catalysed nucleation, even under inhibition. As the '$i$'-state represents the conformation with more $\beta$-sheet content than the dissolved state, we set its interaction strength at $\epsilon_{\text{ii}} = 16$ kT. This high value promotes the conversion of a dissolved oligomer to an intermediate oligomer as well as ensures the stability of the detaching '$i$'-oligomer. The strength of the interaction between monomers in '$i$'-state and '$s$'-state is set to $\epsilon_{\text{si}} = 8$ kT which is in between $\epsilon_{\text{ss}}$ and $\epsilon_{\text{ii}}$. The

interaction between two $\beta$-particles is the strongest interaction in the model: $\epsilon_{\beta\beta} = 60\,\mathrm{kT}$. This high number ensures that the $\beta$-nucleus is well defined as a thermodynamically stable $\beta$-dimer, leaving no ambiguity in the measurements of nucleation rate. The cross-interaction strengths between the dissolved or intermediate and the $\beta$-state are $\epsilon_{\mathrm{s}\beta} = \epsilon_{\mathrm{ss}} + 1\,\mathrm{kT}$ and $\epsilon_{\mathrm{i}\beta} = \epsilon_{\mathrm{ii}} + 1\,\mathrm{kT}$. This means nucleation can happen only in large oligomers where the conversion free energy penalty of changing a single particle within the oligomer to a state with higher $\beta$-content ('$i$'-state) can only be overcome if many new hydrophobic bonds are formed at the same time.

The strength of adsorption $\epsilon_{\mathrm{sf}}$ that promotes binding to the particles that comprise the fibril surface has also been carefully chosen. If this value is too low ($4\,\mathrm{kT}$) there is negligible adsorption to the fibril and no surface-catalysis. But if too high (e.g. $10\,\mathrm{kT}$) there is no detachment of oligomer from the surface and therefore no geometrical rearrangement that allows for conversion to $\beta$-state. At certain conditions [10], the maximum rate of nucleation is achieved at $\epsilon_{\mathrm{sf}}^{max} = 8\,\mathrm{kT}$. We opt for the lower value $\epsilon_{\mathrm{sf}} = 6\,\mathrm{kT}$ in order to avoid possible non-monotonic behaviour of nucleation rate when we change parameters such as chemical potential of the dissolved state and inhibitor properties. The interaction between '$i$'-state monomers and fibril particles is set to $\epsilon_{\mathrm{if}} = 1\,\mathrm{kT}$. This low value promotes the detachment of an '$i$'-oligomer from the fibril while hindering the first conversion of a monomer from dissolved to intermediate state. We vary the inhibitor binding strength in the interval $\epsilon_{\mathrm{If}} = 6 - 8\,\mathrm{kT}$, mimicking the experimental situation where inhibitors have a larger affinity for the fibril surface than unfolded proteins [6]. We nominally set both the inter-inhibitor binding as well as the interaction strength between inhibitors and monomers to zero. The inhibitors interact with other inhibitors or dissolved monomers only via volume-exclusion. Finally, we set the size of the adsorption patch to $l_{\mathrm{a}} = 0.7\,l$ that can accommodate oligomers of large sizes.

**3.3. Monte-Carlo simulation.** Simulations of our coarse-grained computer model were performed using the Monte-Carlo method in a cubic box with periodic boundary conditions and mimicking by the semi-grand canonical ensemble constraints where we kept the volume of the box $V$, the temperature $T$, the chemical potential of monomers $\mu_m$, and the total number of inhibitors $N_I$ constant. Such a scheme was chosen to avoid the depletion of monomers and inhibitors from solution due to adsorption onto the fibril surface and to have good control over the number of particles simulated. All simulations start with a simulation box of size $150w \times 150w \times 150w$, a preformed fibril at the center of the box, 600 randomly distributed dissolved monomers, and $N_{\mathrm{I}}$ number of randomly distributed inhibitors. The preformed fibril contains 92 tightly bound $\beta$-particles and is capped at both ends (unable to grow by elongation) as we want to keep the amount of binding sites constant. We then scale the simulation box to match the specified chemical potential of the dissolved monomers $\mu_m$ by making grand-canonical exchange moves that add or remove dissolved monomers from anywhere in the simulation box, excluding the $D = 12w$ wide exclusion zone around the preformed fibril. During scaling, inhibitors are reinserted whenever they leave the simulation box to keep their number constant.

After initialisation we perform an 'equilibration simulation' where dissolved proteins and inhibitors are allowed to adsorb to the fibril. We run this simulation until we reach chemical equilibrium. We than enable conversion between protein states. During conversion, the position of the particle and the orientation of the spherocylinder's long axis are conserved. But if the peptide changes conformation to a $\beta$-state, we also randomly assign the orientation of the $\beta$-sheet side-patch. The probability of attempting any conformation change was set to $p_{\mathrm{swap}} = 1/5000$, which mimics the slow conversion of the dissolved unfolded protein into a $\beta$-sheet prone configuration.

Each Monte Carlo step involves $N$ translational and orientational moves, where $N$ is the total number of particles in the system (inhibitors, monomers in various states, and fibril particles). Additionally, for $\beta$-state monomers and fibril particles there is an additional random rotation around the spherocylinder's long axis. The chance of these translational and orientational moves being accepted is governed by the Metropolis algorithm. Alternatively, with a chance of $p_{\mathrm{exchange}} = 1/30.000$ in each step, we perform $N_s$ grand-canonical exchange moves, where $N_s$ is the number of dissolved monomers outside the exclusion zone. The simulation terminates when we register two mutually bound $\beta$-particles that define a $\beta$-nucleus.

### 4. Materials and Methods

**4.1. AB42 expression and purification.** The $A\beta_{42}(M1-42)$ peptide comprising residues 671-712 of human APP, MDAEFRHDSGYEVHHQKLVFFAEDVGSNKGAIIGLMVGGVVIA, here called $A\beta_{42}$, was expressed in *Escherichia coli* BL21 DE3 pLysS star from a synthetic gene with *E. coli*-preferred codons in a PetSac plasmid ([13]) in LB medium (10 g/L NaCl, 10 g/L tryptone, 5 g/L Bacto yeast extract, 50 mg/L ampicillin, 30 mg/L chloramphenicol). The peptide was purified as described in [14] using repeated sonication and pelleting of inclusion bodies, followed by dissolution in buffer A (10 mM Tris/HCl, 1 mM EDTA, pH 8.5) with 8 M urea, ion exchange chromatography in batch format using a DEAE cellulose resin with loading in buffer A with 2 M urea, washing in buffer A and elution in buffer A with 50 mM NaCl, and two rounds of size exclusion chromatography in 20 mM sodium phosphate, 0.2 mM EDTA, pH 8.5 on a 26x600 mm Superdex75 column. The purified monomer was lyophilized as multiple identical aliquots.

**4.2. Brichos expression and purification.** Human pro-SPC Brichos was expressed in fusion with thioredoxin-His6 in *E. coli* Origami (DE3) pLysS and purified using metal-affinity chromatography on Ni-NTA-agarose column, cleavage by thrombin, a second metal-affinity chromatography step to remove the thoredoxin-His6 tag and finally ion exchange chromatography, as described previously [15, 16].

**4.3. AB42 kinetics in the presence of Brichos.** Just prior to the experiments, an aliquot of purified $A\beta_{42}$ monomer was dissolved in 1 mL 6 M GuHCl, 20 mM sodium phosphate, pH 8.5 and monomers again isolated using size exclusion chromatography in 20 mM sodium phosphate, 0.2 mM EDTA, pH 8.0 on a 10x300 mm Superdex75 column. The peptide was collected in a low-binding tube (Axygen) on ice, concentration determined from the integrated absorbance at 280 nm of the collected fraction (using $\epsilon_{280} = 1440\,\mathrm{l\,mol^{-1}cm^{-1}}$). A sample series containing proSPC-Brichos at varying concentration (0-3 µM) and the following components at constant concentration: 3 µM A42 monomer, 30 nM A42 seed, 10 µM thioflavin T (ThT), 20 mM sodium phosphate, 0.2 mM EDTA, pH 8.0 was prepared in low-binding tube (Axygen) on ice. The solutions were transferred to multiple wells of a 96-well plate (PEG-ylated black polystyrene with clear bottom, Corning 3881) and placed in a plate reader (BMG optima) thermostated to 21 °C (placed in a cold cabinet to allow thermostating at such temperature). The ThT fluorescence was read through the bottom of the plate under quiescent conditions with excitation at 440 nm and emission at 480 nm. The seed was collected when reaching the plateau in a previous reaction of A42 alone monitored by ThT fluorescence in the same manner.

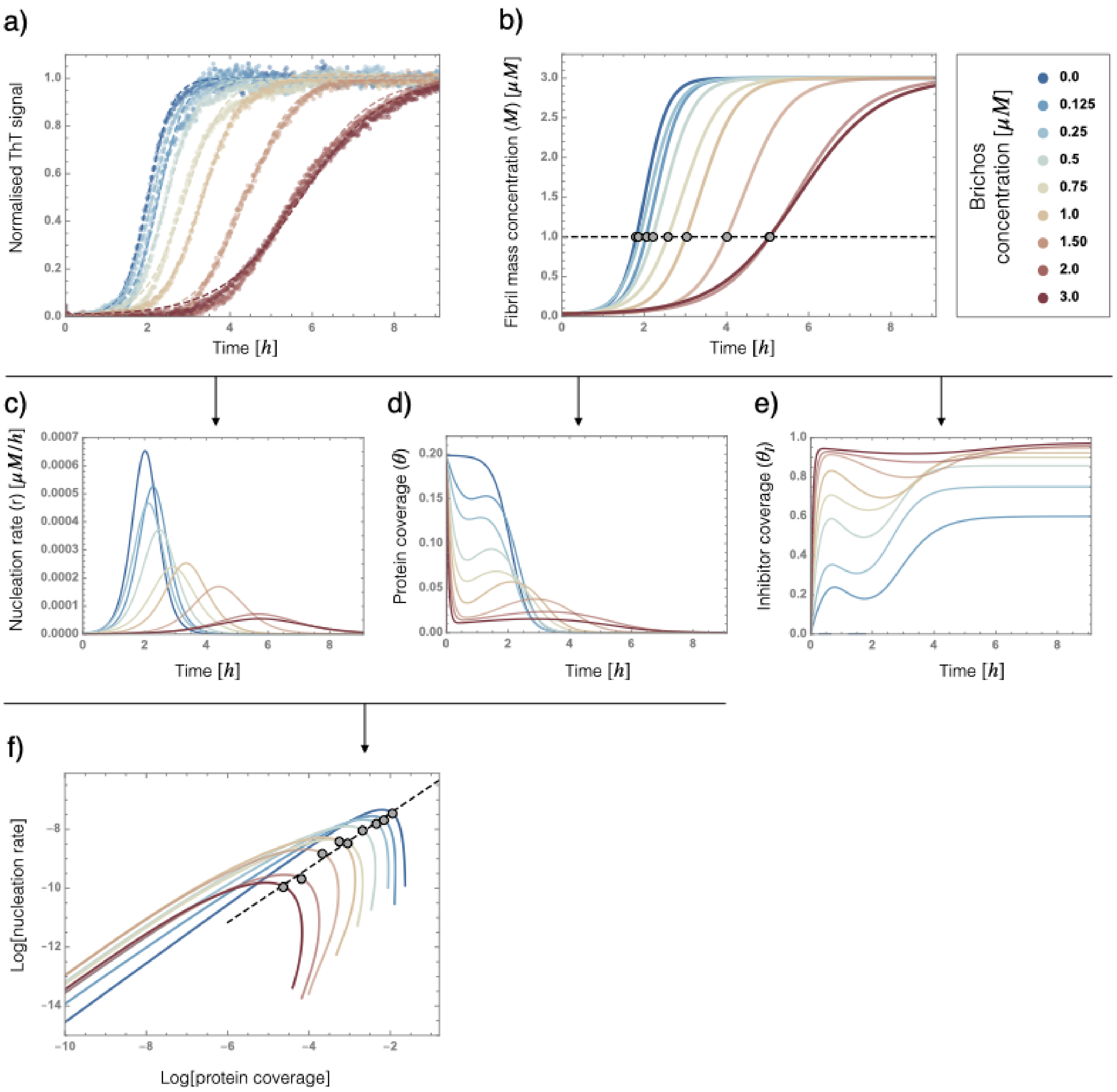


**Fig. S1. Experimental analysis workflow to extract $\sigma_{\mathrm{I}}$.** a) Normalised data for A$\beta_{42}$ aggregation as measured by ThT fluorescence at different concentrations of proSP-C Brichos chaperone. Experiments were performed at temperature $T = 21^{\circ}C$ with initial A$\beta$ concentration of $m_0 = 3\mu M$, and a small amount of preformed seed $M = 0.03\mu M$. We fit the data at each Brichos concentration with a closely fitting sigmoid to get an analytical and differentiable representation of data. b) Fibril mass concentration $(M)$ as a function of time. Slicing the dataset with a horizontal line gives us times at which a certain amount of protein is aggregated to fibril form for each concentration. For example, grey circles track the times at which $1\mu M$ of protein is aggregated. From the normalised data we can extract the rate of nucleation (c) using Eq. S.3 and the binding kinetics for protein (d) and inhibitor (e) coverage, based on Eqs. S.6, S.7. f) We plot the nucleation rate against protein coverage as a parametric plot on a log-log scale and by fixing the fibril mass for each concentration – for example $M = 1\mu M$; grey points in f) correspond to grey points in b) – we get the scaling $\sigma_{\mathrm{I}}$.

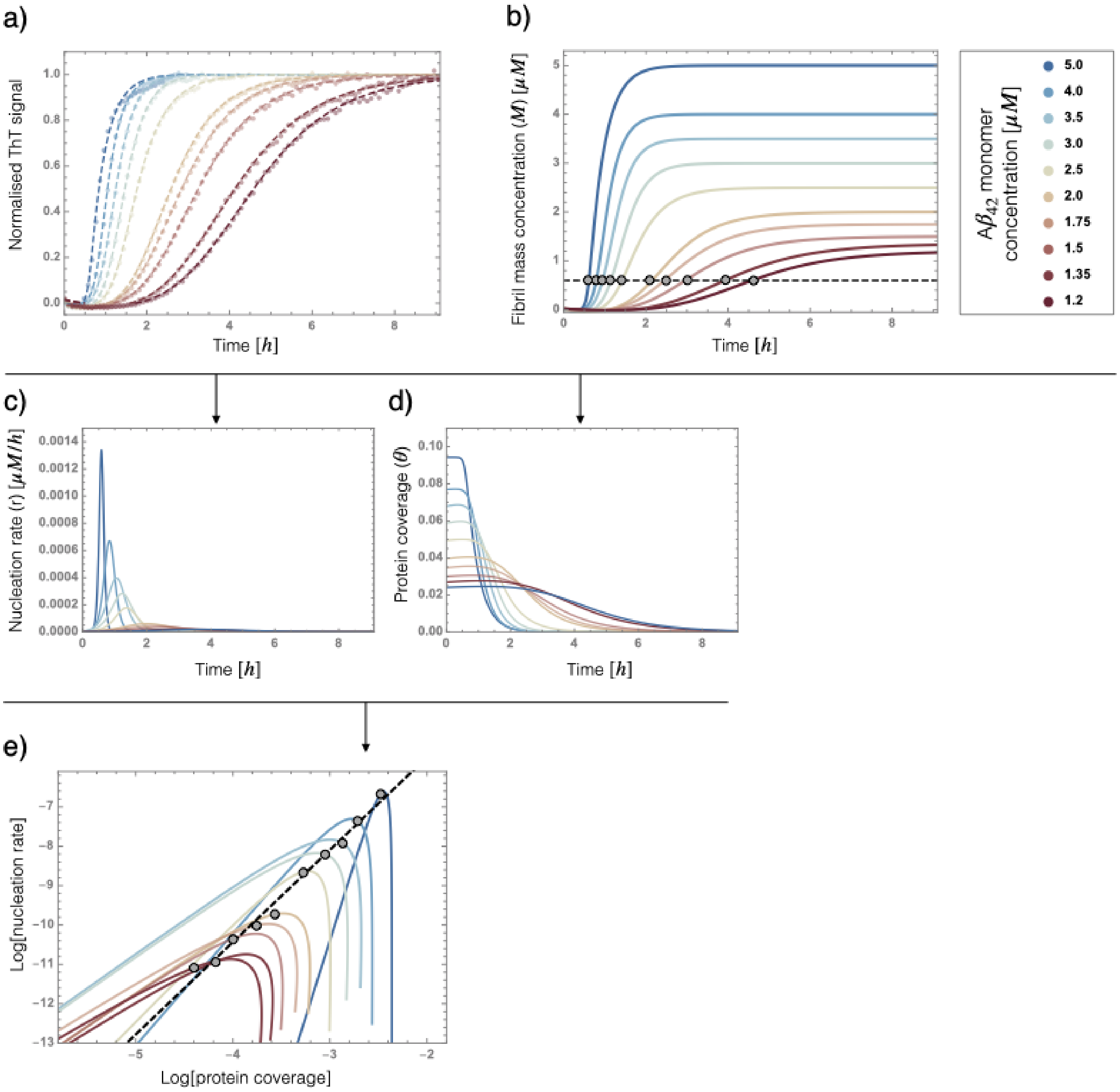


**Fig. S2. Experimental analysis workflow to extract** $\sigma$. a) Normalised data for A$\beta_{42}$ aggregation as measured by ThT fluorescence at different concentrations initial monomer concentration. Experiments were performed at temperature $T = 37°C$ without any inhibitor or fibril seed. We fit the data at each concentration with a closely fitting sigmoid to get an analytical and differentiable representation of data. b) Fibril mass concentration $(M)$ as a function of time. Slicing the dataset with a horizontal line gives us times at which a certain amount of protein is aggregated to fibril form for each concentration. For example, grey circles track the times at which $0.6\mu M$ of protein is aggregated. From the normalised data we can extract the rate of nucleation (c) using Eq. S.3 and the binding kinetics for protein (d) coverage, based on Eq. S.6 (for $\theta_I = 0$). f) We plot the nucleation rate against protein coverage as a parametric plot on a log-log scale and by fixing the fibril mass for each concentration – for example $M = 0.6\mu M$; grey points in f) correspond to grey points in b) – we get the scaling $\sigma$.

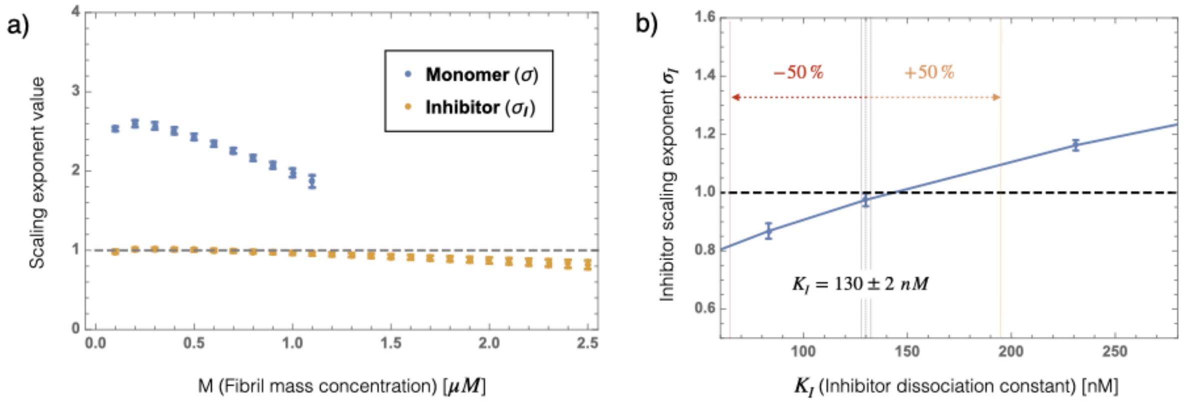


**Fig. S3. Dependence of scaling exponents**. a) Value of scaling exponents $\sigma$ (blue data), and $\sigma_\mathrm{I}$ (orange data) at different amounts of catalytic fibril surface, as given by the concentration of aggregated fibril mass. The error corresponds to the uncertainty in the slope of the linear fits from Fig. S1f (for $\sigma_\mathrm{I}$), and from Fig. S1e (for $\sigma$). b) Sensitivity analysis on how the inhibition scaling exponent depends on the values of inhibitor binding constants. At the literature value for the inhibitor dissociation constant $K_\mathrm{I} = (130 \pm 2)\mathrm{nM}$ we find $\sigma_\mathrm{I} = 0.98 \pm 0.03$. Artificially increasing the uncertainty in $K_\mathrm{I}$ to $50\%$, we find the uncertainty in value of $\sigma_\mathrm{I}$ increases to $\sigma_\mathrm{I} = 1.0 \pm 0.2$.

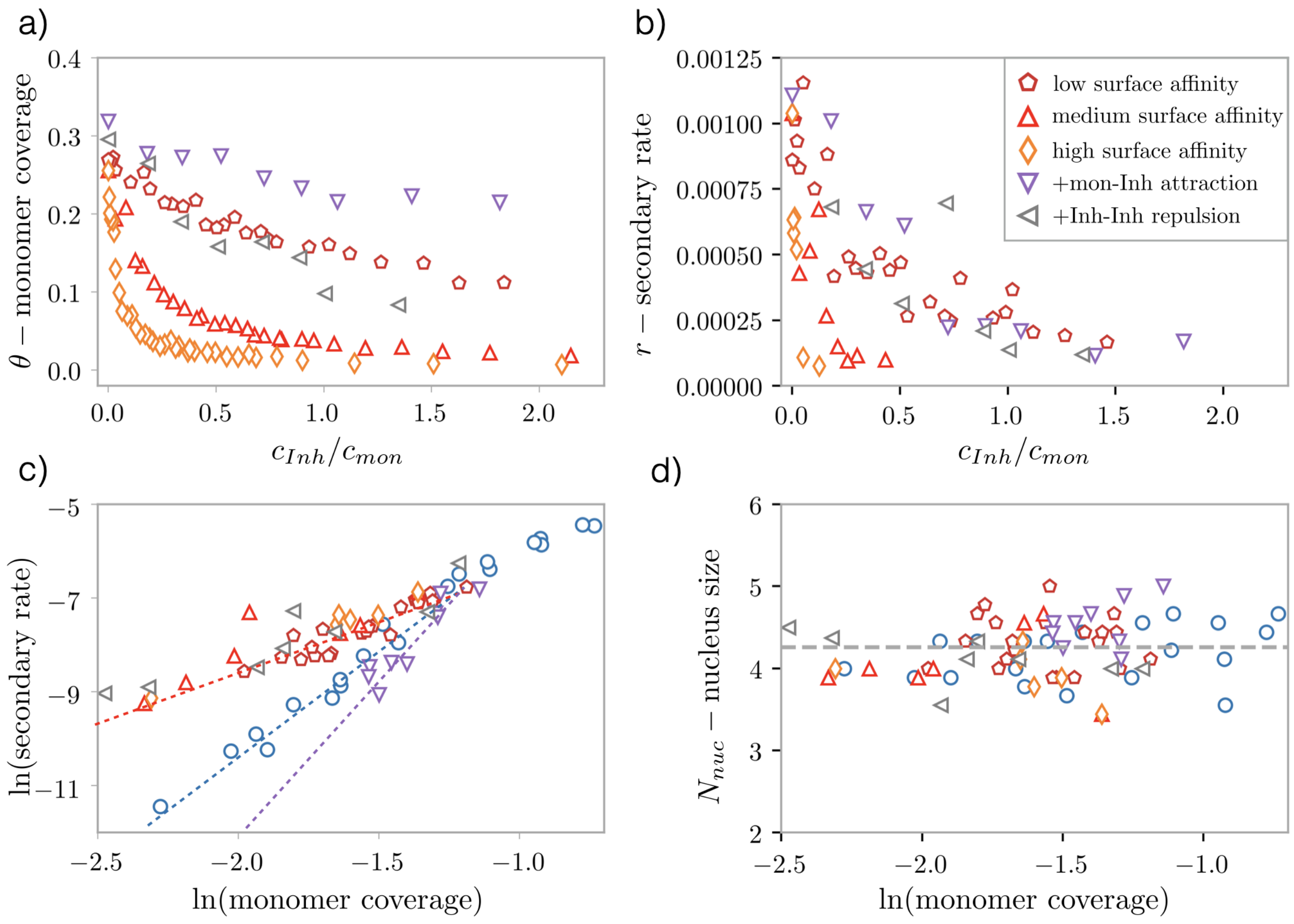


**Fig. S4. Results from coarse-grained computer simulations of secondary nucleation and its inhibition** a) Keeping monomer concentration constant we look at the effect on protein coverage when adding inhibitor into solution. Dark red pentagons, red upward triangles and orange diamond represent data for inhibitors with only volume-exclusion interaction with other particles in the system but with increasing fibril affinities ($\epsilon_{\mathrm{If}} = 6kT$, $\epsilon_{\mathrm{If}} = 7.2kT$, and $\epsilon_{\mathrm{If}} = 8kT$, respectively). Violet downward triangles and grey leftward triangles in addition present data with inhibitors that have an tip-to-tip attraction of $\epsilon_{\mathrm{Is}} = -4kT$ between inhibitors and dissolved monomers, and inhibitors with tip-to-tip repulsion of $\epsilon_{\mathrm{II}} = 4kT$, respectively. In all cases we find that protein coverage drops with inhibitor concentration due to competitive binding but to a different degree. b) The rate of secondary nucleation also drops with inhibitor concentration for all types of inhibitors. c) To measure the scaling exponents we plot the rate against coverage in log-log scale in the absence of inhibitor (blue circles, changing monomer concentration) and at presence of different types of inhibitors. For only volume-exclusion interactions between inhibitors and monomers we find $\sigma_{\mathrm{I}} < \sigma$ and all data falls on the same line, indicating that affinity to the fibril surface and the interactions between inhibitors do not influence $\sigma_{\mathrm{I}}$. In the case of attractive monomer-inhibitor interactions we get $\sigma_{\mathrm{I}} > \sigma$. d) Finally, we show that the size of the nucleating species is the same in the absence and presence of different types of inhibitors at all protein coverages. This indicates that inhibitors only influence the protein oligomer distribution but not also the conversion step.

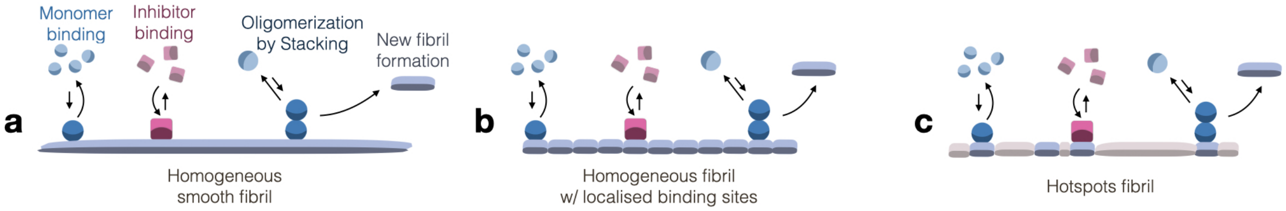


**Fig. S5. Dependence of scaling exponents**. Schematic of the secondary nucleation mechanism and its inhibition where Ab42 oligomers are formed by stacking of monomers on top of each other.
